# A cell-free strategy for profiling intracellular antibiotic sensitivity and resistance

**DOI:** 10.1101/2023.04.13.536698

**Authors:** Kameshwari Chengan, Charlotte Hind, Lakshmeesha Nagappa, Matthew E. Wand, Tanith Hanson, Ruben Martin Escolano, Anastasios Tsaousis, José A Bengoechea, J. Mark Sutton, Christopher M Smales, Simon J Moore

## Abstract

Antimicrobial resistance (AMR) is a pandemic spread across multiple priority infectious disease threats. While the cell envelope plays a key role in AMR, this also makes it challenging to study how antibiotics function inside the cell. Herein, we present a *Klebsiella pneumoniae* cell-free gene expression (CFE) platform for the rapid profiling of intracellular antibiotic sensitivity and resistance. This cell-free approach provides the unique macromolecular and metabolite components from this microbe, which include multiple antibiotic targets from transcription, translation, and metabolic processes. First, we compare the *K. pneumoniae* CFE system to whole cell antimicrobial assays. We find that several antibiotic classes show higher sensitivity in the CFE system, suggesting limitations in antibiotic transport in the whole cell assay. Next, we evolved *K. pneumoniae* strains with resistance to specific antibiotics and use whole genome sequencing analysis for genotyping. As an exemplary case, we show that a single RNA polymerase beta subunit variant H526L (also frequently found in multidrug resistant *Mycobacterium tuberculosis*) confers a 58-fold increase in CFE resistance to rifampicin. Overall, we describe a safe (i.e., non-living, non-pathogenic) platform suitable for studying an infectious disease model in a Containment Level 1 laboratory. Our CFE strategy is generalisable to laboratory and clinical *K. pneumoniae* strains and provides a new experimental tool to profile intracellular AMR variants. In conclusion, our CFE tool provides a significant advance towards understanding AMR and complements wider infectious disease studies.

## Introduction

*Klebsiella pneumoniae* (and *K. oxytoca*), is a leading cause of hospital-acquired bacteriaemia deaths, which carry a mortality rate of up to 41% after infection with carbapenem-resistant strains^1^. Like other Gram-negative bacterial pathogens, *K. pneumoniae* presents a “phalanx” of cell envelope defences that act as a barrier against many antibiotic classes^2^. This includes an outer and inner cell membrane with orthogonal entry properties, peptidoglycan cell wall, multiple antibiotic efflux transporters, and an extracellular capsular polysaccharide layer^3^. The capsule is also a key virulence factor^4–6^. Considering these physical barriers, it remains difficult to study the mechanisms of AMR in *K. pneumoniae*, especially those in the cytoplasm and intracellular macromolecular systems. Therefore, there is an urgent need to develop novel approaches to address this problem, to facilitate the development and identification of new antimicrobials and approaches that prevent or circumvent AMR.

To monitor the spread of AMR, there are two general approaches: genotyping and phenotyping. Genotyping uses whole-genome next-generation sequencing (NGS) and bioinformatics tools, such as RESfinder^7, 8^, to detect genetic determinants associated with AMR (i.e., resistance genes, single-nucleotide polymorphisms in AMR determinants). Uniquely, NGS can study single strains and mixed populations. However, both genotyping and bioinformatics analysis is reliant upon accurate laboratory-based phenotypic experiments. This involves cell culture, broth dilution, time kill, and disc diffusion assays, following the recommended standards from the U.S. Food and Drug Administration (FDA), World Health Organization, National Committee for Clinical Laboratory Standards and EUCAST. At the phenotyping level, limitations exist in elucidating the precise mechanisms that underpin AMR within infectious diseases such as *K. pneumoniae*. First, antimicrobial inhibition of whole cells is measured by a minimum inhibitory concentration (MIC) value. MIC_90_ and MIC_50_ values are the lowest concentration of antibiotic that inhibit 90 and 50% of isolates, respectively. Critically, the MIC_50_/_90_ value is a complex measure of overall antibiotic activity, a combination of antibiotic import, efflux, metabolism, selectivity, and potency, all overlaid by mutations associated with the resistome. Additionally, mutations associated with a specific AMR phenotype, can confer collateral resistance or sensitivity to structurally or functionally unrelated antibiotics^9–11^. Similarities occur in cancer drug resistance^12^. To study single AMR mechanisms in isolation, this generally requires *in vitro* studies of purified proteins or complexes (e.g., ribosomes) through specialist biochemistry and structural biology approaches. In addition, isolated single targets lack the associate macromolecular components and interactions critical to core processes such as gene expression.

In considering an alternative approach, *in vitro* translation (IVT) systems are a proven tool for measuring ribosomal inhibitors for mode of action studies^13, 14^. IVT systems require a cell extract, RNA template, amino acids, nucleotides, and a metabolite mixture to catalyse protein synthesis in a ‘one-pot’ biochemical reaction. IVT and cell-free systems lack a cell membrane and genome. While this is a potential caveat of the system, it also provides many advantages in that by removing antibiotic transport and genetic regulation, IVT systems provide a direct surrogate model to study antimicrobials that disrupt gene expression. In addition, IVT permits high-throughput screening (HTS). For example, HTS with an *E. coli* IVT system screen of 20,000 natural product extracts identified a novel tetrapeptide (GE81112) ribosomal inhibitor^15^, which disrupts fMet-tRNA interaction with the 30S ribosomal subunit. However, most antimicrobial studies with cell-free systems are limited to laboratory adapted models such as *Escherichia coli* and *Bacillus subtilis*^13, 14^. An exception is a *Streptococcus pneumoniae in vitro* transcription-translation (IVTT) system, which discovered a naphthyridone ribosomal inhibitor (A-72310)^16, 17^, which also separately inhibits DNA gyrase (GyrA) in Gram-negative bacteria^18^.

Recently, we and others have developed a wide range of cell-free gene expression systems (CFE) – also referred to as cell-free protein synthesis (CFPS) – from both Gram-negative and Gram-positive bacterial cell types^19–24^. This has enabled an alternative approach to investigate established areas of research. Together with optimised *E. coli* CFE systems^25–29^, non-model CFE systems carry additional functionality such as the native macromolecular machinery and metabolic enzymes unique to these microbes. Specifically, bacterial cell extracts typically contain a few hundred proteins^20, 27^ from processes such as protein synthesis^28, 30^ and transcription^31^, while there are a plethora of enzymes from central metabolism^24, 32–35^. While *E. coli* is the most studied, it is known that this system harbours an active electron transport chain and ATP synthase complex, contained within inverted membrane vesicles that form during cell lysis^32^. Together with active central metabolic pathways, CFE reactions catalyse ATP regeneration and accumulate inhibitory waste products (e.g., lactate, acetate)^36^. Underlying all these intracellular processes is a myriad of background protein-protein interactions (PPIs), a potential novel target for antimicrobial development^37, 38^. Therefore, CFE systems provide a unique opportunity for studying antimicrobials, since there is a potential array of different antibiotic targets – unique to each microbe – that impact protein synthesis (and cell fitness), either through inhibition of transcription, translation, metabolism, or PPIs.

In this study, we describe the development of a CFE system from *K. pneumoniae*, a major World Health Organisation (WHO) priority pathogen, allowing the study of antibiotic target susceptibility and resistance in a safe (non-pathogenic) surrogate model. Here we apply drug susceptibility profiling, evolutionary and genomics studies to explore the relationship of cell and cell-free antibiotic resistance in a range of *K. pneumoniae* laboratory and clinical strains. To start, we optimise a *K. pneumoniae* CFE model platform, developed from the ATCC 13882 quality control strain, which we apply to screen known antimicrobials and compare their activity to cell-based growth assays. From this, we reveal key differences in antibiotic potency ascribable to uptake/efflux barriers in cell-based assays. We also extend this CFE platform to include systems derived from multidrug-resistant strains. This includes two clinical strains (ST258-T1b and NJST258-1) and three isogenic mutants (derived from ATCC 13822 strain) with elevated resistance to single antibiotics. Finally, we deploy the CFE platform to study how single AMR genetic determinants influence gene expression. This provides significant advantages, since the removal of the cell envelope and genomic DNA allows us to study antimicrobial targets within the context of a complete cytoplasmic milieux, thus providing a robust model, both for drug discovery and studying AMR.

## Results

### Establishing a rapid and safe *K. pneumoniae* CFE platform from laboratory and clinical isolates

A range of CFE systems have recently emerged from diverse bacterial genera, providing a fresh approach to study non-model microbes^19–24^. To make a highly active CFE system, this requires the isolation of a cell extract enriched with active ribosomes from rapidly dividing cells. Then, a gene expression reporter system (i.e., plasmid DNA) and a metabolite solution (i.e., amino acids, nucleotides, energy source and cofactors) is added to the cell extract to initiate coupled transcription-translation (**Figure 1a**). To generate a *K. pneumoniae* CFE system, we selected the ATCC 13882 standard strain used for quality control in antimicrobial susceptibility testing^39^. We then optimised the growth of *K. pneumoniae* ATCC 13882 (**Figure S1-3**) and processing steps to provide concentrated cell extracts (20-24 mg/mL). Initially, we tested a range of standard plasmids, including Pr-deGFP-MGapt (**Table S1**), which is highly active in *E. coli* CFE systems, since *K. pneumoniae* is not too distantly related. However, we were unable to detect protein synthesis. Instead, we tested the pTU1-A-SP44-*mScarlet-I* (pSJM1174) plasmid that is optimised for *Streptomyces* gene expression. This plasmid contains a constitutive promoter driving the transcription of the encoded mScarlet-I red fluorescence protein. With some sequential optimisation (**Figure S4-S10**), mScarlet-I synthesis reached up to 5.73 ± 1.02 µM (**Figure 1b**). Further fine tuning of the reaction conditions could afford yields of up to 11 µM mScarlet (**Figure S11**). While protein synthesis rates were modest (∼0.5 aa/s) compared to *E. coli* CFE (>1.5 aa/s^27^), the reactions were active for up to 14-16 hours; new CFE systems are typically only active for ∼2-6 hours. Next, we considered the safety level of *K. pneumoniae*, especially for the clinical isolates, which requires handling in Containment Level 2 (CL2) facilities. CFE systems are non-living^27^ and remain stable at -80°C^40^. To confirm this, we tested the cell extracts for colony forming units (CFU). We found approximately 10^2^ CFUs per mL of concentrated cell extract (**Figure S12**). However, these viable cells were easily removed by passing the cell extracts (20 mg/mL) through a 0.2 µm filter without altering CFE activity (**Figure S13**). Removal of viable cells allows transfer of cell extracts from a CL2 to a CL1 lab. We also confirm that the cell extracts remain stable and active for up to 4 years during our study, when stored at -80°C (data not shown).

**Figure 1.**
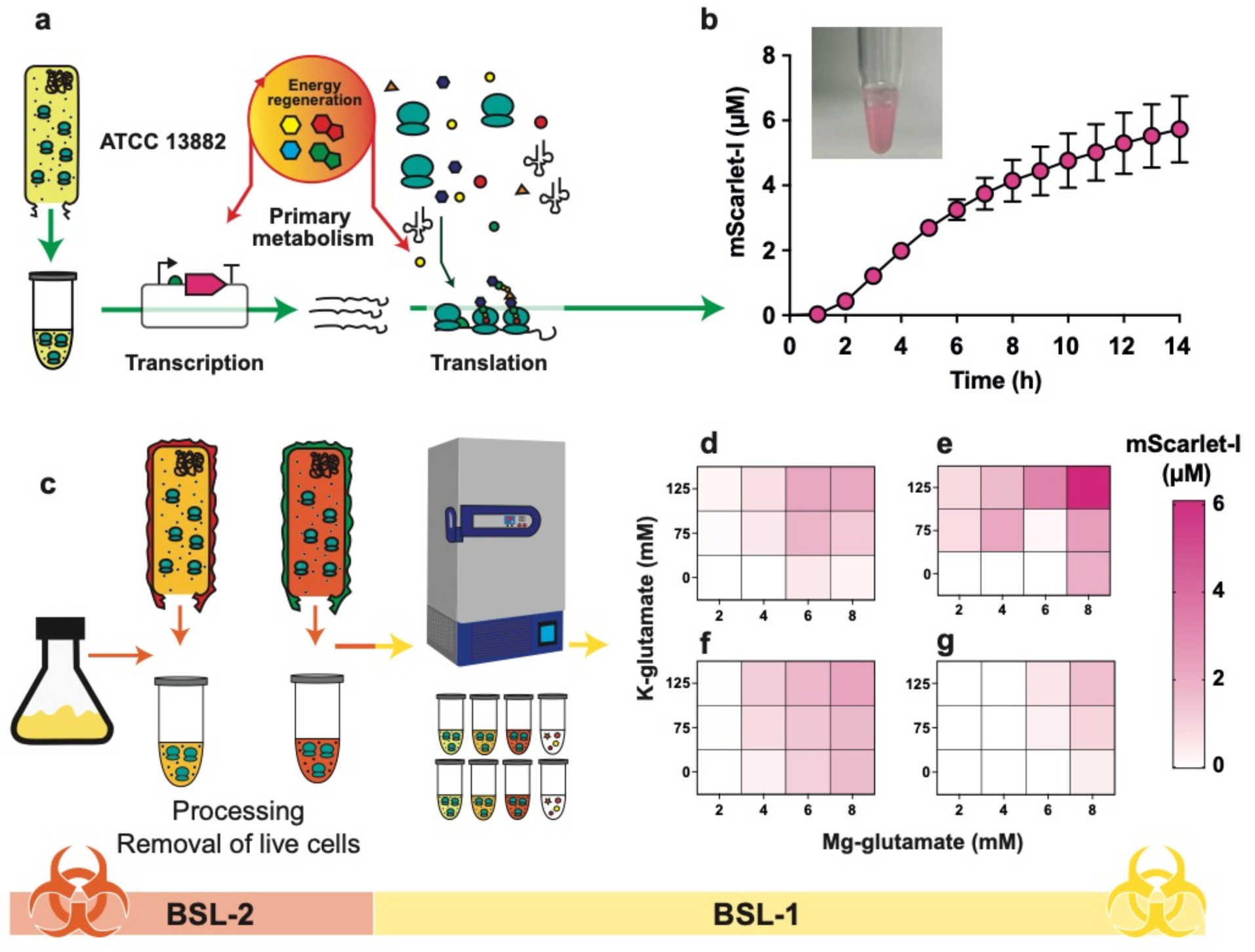
A *K. pneumoniae* CFE platform from laboratory and clinical isolates. (**a**) Overview of components and stages of the CFE reaction. (**b**) *K. pneumoniae* ATCC 13882 CFE activity of batch reactions (visual images) and real-time measurement of mScarlet-I (μM) concentration (calculated from a reference standard) over 14 hours. Data is mean and standard deviation of two biological repeats. (**c**) Workflow for CFE extract processing of clinical strains. (**d-e**) A heatmap representing the optimisation of mScarlet-I (μM) synthesis from ST258-T1b and (**f-g**) NJST258-1 clinical CFE systems with Mg-glutamate and K-glutamate. Two independent repeats (left and right panels) are shown from four biological repeats.

Next, to show flexibility in the CFE approach to study other strains, we repeated the process with the multi-antibiotic resistant ST258-T1b and NJST258-1 clinical isolates (**Figure 1c**). We obtained reasonable CFE activity for these clinical strains by optimising the magnesium-glutamate and potassium-glutamate levels. Magnesium is required for bioactivity of ATPases and ribosomes. Glutamate provides additional energy through the Krebs cycle via α-ketoglutarate^41^. This essential metabolite is highly abundant (∼96 mM) inside *E. coli* cells, and is a key source of energy and nitrogen^42^. In comparison to the ATCC 13882 strain, NJST258-1 and ST258-T1b CFE systems produced less mScarlet at ∼2 µM and ∼4.5 μM (*n* = 2 biological repeats), respectively (**Figure 1d-g**). At this point, we initially attempted to explore antibiotic-resistance in these clinically relevant CFE systems. However, we encountered issues with variability between extract batches (**Figure S14**), likely due to silent gene expression or loss of plasmid-born resistance elements in the preparation of cells for extract processing. Instead, we chose to focus on characterising the ATCC 13882 strain, due to its superior activity and minimal batch variation, which we used to profile known antimicrobials, while retaining a view to using this platform later to study antibiotic resistance with CFE systems.

### High sensitivity and selectivity of the *K. pneumoniae* CFE system for inhibitors of transcription and translation

Next, we aimed to profile the *K. pneumoniae* ATCC 13882 CFE system for antimicrobial selectivity and sensitivity. This was performed following recommended standards for drug screening^43^, where we considered the need for a strong signal-to-background (S:B) ratio and statistical robustness. First, we scaled-up extract preparation and optimised the magnesium and potassium glutamate concentration. We discarded extract batches with less than 50% of the maximum recorded activity (11.0 ± 0.73 µM) to standardise the robustness of the assay. Measured from two biological repeats, this data provides a strong S:B ratio of 134:1 between the error margins (**Figure 1b**). To validate the specificity and sensitivity, we next tested a selection of control antibiotics. This included kanamycin and erythromycin, both ribosomal inhibitors. Ampicillin was also included as a non-specific inhibitor, since its target is cell wall biosynthesis, which should not affect CFE activity. For the clinical antibiotics with intracellular targets, estimated IC_50_ values were within the expected range – i.e., low micromolar (**Figure S15**). The CFE assay was insensitive to ampicillin up to 100 μM, the highest concentration tested within the model. To validate the experimental approach, a robust Z’ score was also determined between 0.72 to 0.88 for eight 384-well microtiter plates (**Figure 2a**) – a minimum robust Z’ score of 0.5 is recommended for a new assay^43^.

**Figure 2.**
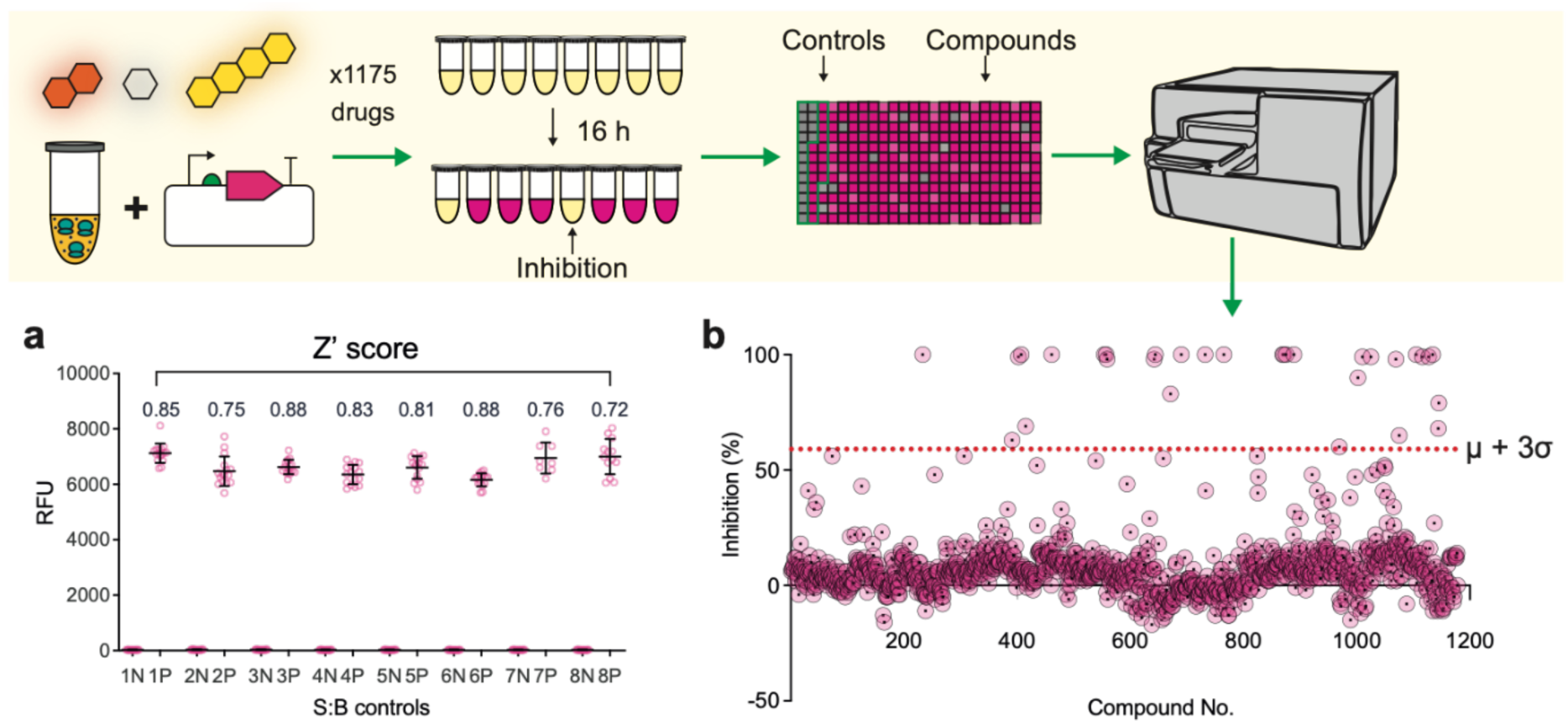
Antimicrobial screening with *K. pneumoniae* ATCC 13882 CFE. (**a**) Z’ scores and fluorescence readings of negative (N) and positive (P) control reactions from eight separate 384-well plates (numbered 1-8 – e.g., 1N, 1P). The mean and standard deviation of the relative fluorescence units (RFU) of 16 independent repeats is shown. (**b**) Final (16 h) measurements of FDA-library drug screen, set up with 1% DMSO and controls, with a standard *K. pneumoniae* ATCC 13882 CFE reaction as outlined in the methods. Each data point is the mean of two technical readings. Data is shown converted to percentage (%) inhibition relative to the mean fluorescence of the positive control reactions containing 1% (v/v) DMSO.

A typical industrial high-throughput screening (HTS) campaign requires enhanced assay stability at room temperature for several hours. The *K. pneumoniae* CFE reaction requires all three components (cell extract, energy solution and DNA) to initiate protein synthesis. We incubated cell extracts with plasmid DNA or energy solution for several hours before addition of the remaining component. Interestingly, the cell extract and DNA mixture remained stable for up to 4 hours at room temperature without significant loss of mScarlet-I yield (**Figure S16)**. In contrast, the cell extract and energy solution showed a time-dependent loss of activity, with a >50% decrease after 2 hours. This suggests the energy solution is the major source of instability due to metabolic activity within the cell extracts, whereas the DNA is stable. Finally, we tested the *K. pneumoniae* CFE system with increasing amounts of dimethylsulphoxide (DMSO), which is required for drug solubility in bioassays. The CFE reaction tolerated up to 2.5% DMSO (v/v) without a significant change in activity (**Figure S17**).

To assess the specificity and selectivity of the *K. pneumoniae* CFE system, we performed an antimicrobial screen against a known library. We used the Selleck FDA-approved drug library (1175 compounds) since it contains most clinically used antibiotics. Initially we performed a preliminary screen at a nominal concentration of 10 μM and 100 μM with the infectious disease sub-set library, which includes antibacterial, antifungal, and antiviral compounds (**Supporting File 1**). At 100 μM, a few non-specific hits (e.g., cell wall / membrane targeting antibiotics) inhibited the *K. pneumoniae* CFE assay. At 10 μM, most transcription or translation targeting antibiotics were detected suggesting this was an appropriate starting concentration. We then screened the FDA library with 1% (v/v) DMSO at this concentration, as outlined in the methods. By applying a sigma threshold (3𝜎 + <u>𝑥</u>), 31 compounds (2.63% of library) were confirmed as positive hits, all known bacterial transcriptional or translational inhibitors (**Figure 2b**). A further 67 compounds were below the sigma threshold but showed ≥20-90% inhibition (mid-range hits). These compounds contained known antibiotics with intracellular targets (e.g., ribosome), as well as non-specific drugs.

Since the primary screen applied an arbitrary concentration of 10 μM, a secondary screen was performed at 100 μM for the 69 compounds that showed inhibition between ≥20-90%. This verified for false negatives below the sigma threshold. Compounds showing ≤30% inhibition remained inactive at higher concentrations, except for chlortetracycline and florfenicol (both ribosomal inhibitors) and teniposide and ethacridine lactate which showed increased activity. Ethacridine (antiseptic) and teniposide (DNA targeting) likely inhibit through a non-specific mechanism at this high concentration. Between ≥30-90% most compounds that had increased or unaltered activity at 100 μM were known antibiotics or antitumour (DNA targeting) agents. These data show that although the sigma threshold was estimated at 59%, there were several known antimicrobials below this threshold. Interestingly, within this mid-range activity, there were several fluoroquinolone analogs and the related novobiocin (target DNA gyrase), which even at 100 μM did not reach 100% inhibition (**Supporting File 1**). Here, it was important to compare the *K. pneumoniae* CFE system to a known model, so we chose *E. coli* MG1655 CFE. We prepared cell extracts from MG1655 as previously described^24^, using the same CFE reaction conditions as *K. pneumoniae* CFE except that *E. coli* MG1655 CFE reactions also required additional aeration to support oxidative phosphorylation. This process appears non-functional in *K. pneumoniae* CFE since the reactions run microaerobic (i.e., static) without a significant decrease in activity. This may relate to differences in the rupture of the cell envelope during cell lysis and the formation of inverted vesicles, which for *E. coli* at least, houses an active electron transport chain and ATP synthase to regenerate energy from a carbon source^36^. After optimising the reaction conditions, the *E. coli* MG1655 CFE system produced up to ∼3 μM of mScarlet-I. To investigate the specificity for both the *K. pneumoniae* and *E. coli* CFE assays, we screened 218 compounds associated with antimicrobial effects. Overall, the *E. coli* MG1655 and *K. pneumoniae* ATCC 13882 CFE systems performed similarly with an r^2^ coefficients of 0.63 and 0.73 at 10 μM and 100 μM drug concentration, respectively (**Figure 3a-b**). However, there was an apparent skew in correlation that indicates *E. coli* CFE is more sensitive to several ribosomal and DNA gyrase (i.e., fluoroquinolones, nalidixic acid) inhibitors than the *K. pneumoniae* CFE system (**Figure 3a-b**). First, there were some known ribosomal inhibitors (e.g., lincomycin, linezolid) that displayed minimal inhibition at 10 μM for *E. coli* and *K. pneumoniae* CFE (**Figure 3a**) but showed >85% inhibition at 100 μM for both systems (**Figure 3b**). In the case of linezolid, this is known to be more effective for Gram-positive bacteria^44^. As expected, there were also several non-specific hits at 100 μM (**Figure 3b**). However, at 10 μM of drug, there were only three non-antibacterial inhibitors above the sigma threshold disrupting both CFE systems. This included moroxydine, caspofungin and triclabendazole. Interestingly, both triclabendazole (flukeworm drug) and caspofungin (anti-fungal) were recently reported to show antibacterial activity^45, 46^. Additionally, there were six non-specific inhibitors (levamisole, camptothecin, terbinafine, artemisinin, cetylpyridinium and piperacillin) that fully inhibited *E. coli* CFE at 10 μM, with only partial inhibition of the *K. pneumoniae* CFE system ranging from 10% to 69%. Surprisingly, piperacillin showed strong activity for both systems, but this was not consistent with other penicillin analogs. In general, the combined compound library screens indicate that *E. coli* CFE was more sensitive to non-specific inhibitors than the *K. pneumoniae* CFE system.

**Figure 3.**
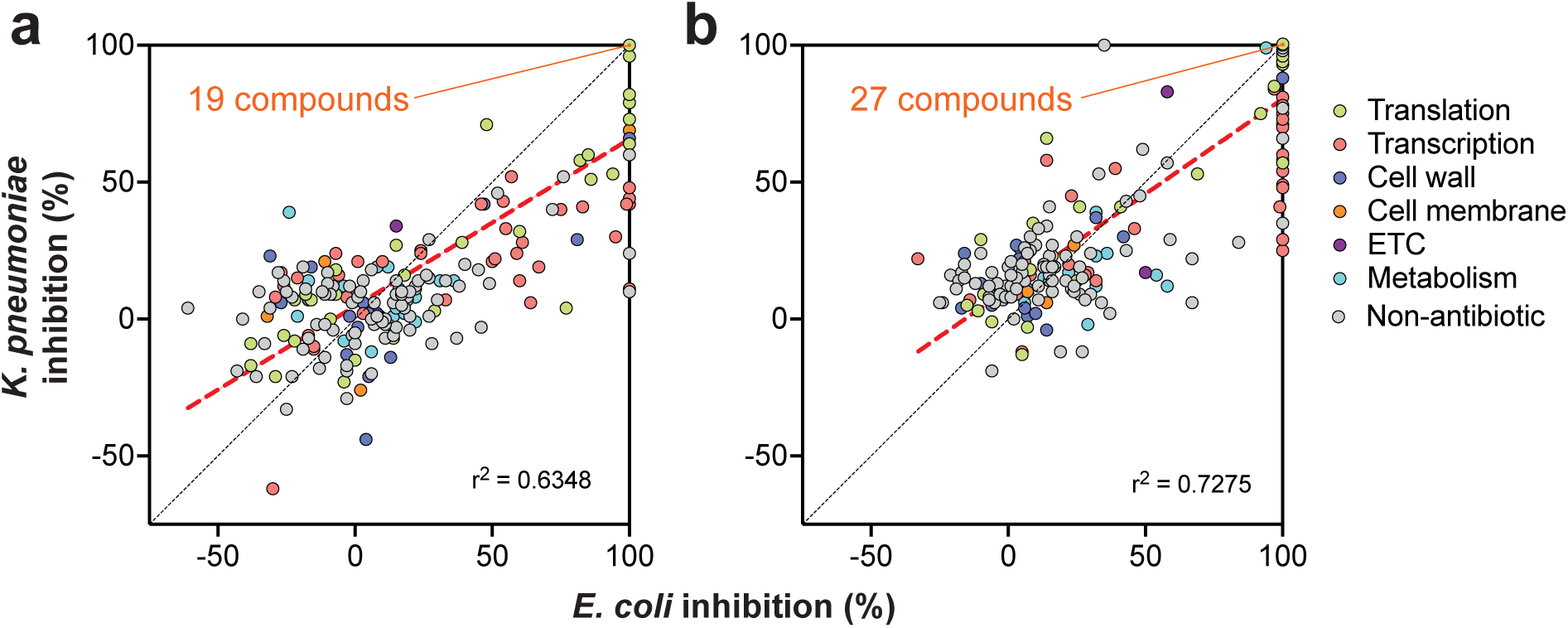
A comparison of *E. coli* and *K. pneumoniae* CFE inhibition by 218 compounds from the Selleck infectious disease subset library. **(a)** 10 μM and **(b)** 100 μM drug concentration. Data points are colour coded to respective biological targets. The “non-antibiotic” category includes anti-viral, ant-fungal, anti-parasitic and anti-helminth inhibitors. Some inhibitors showed 100% inhibition for both *E. coli* and *K. pneumoniae* CFE, which is indicated within the graph as an orange line and text as a total number. Data is represented as normalised inhibition (%) relative to the experimental controls for two technical repeats. A line of best fit is plotted as a red dashed line, with corresponding r^2^ value indicated bottom right. A line of identity is plotted as a black dashed line.

A surprising finding was that almost all the fluoroquinolone analogs fully inhibited *E. coli* CFE, but not *K. pneumoniae* CFE, complementing our earlier observations (**Figure 4**). Fluoroquinolones inhibit DNA gyrase that leads to dsDNA breaks. We separately tested a selection of these compounds for activity in *K. pneumoniae* ATCC 13882 whole cell assays and observed low micromolar MIC values for all compounds (**Table S2**). This suggests inhibition of DNA gyrase only partially inhibits the *K. pneumoniae* CFE reaction, possibly because of biochemical differences within the CFE system during unwinding of supercoiled DNA. While this effect is lethal to a cell (i.e., dsDNA breaks), it has a relatively modest impact on the *K. pneumoniae* CFE system. The mechanism by which it only partially reduces transcription-translation remains unclear at this stage but identifies a unique difference and limitation in comparison to *E. coli* CFE. The difference in activities for the fluoroquinolone and nalidixic acid analogs was intriguing but suggests the *K. pneumoniae* CFE assay is best suited to studying antibiotics that directly target transcription or translation.

**Figure 4.**
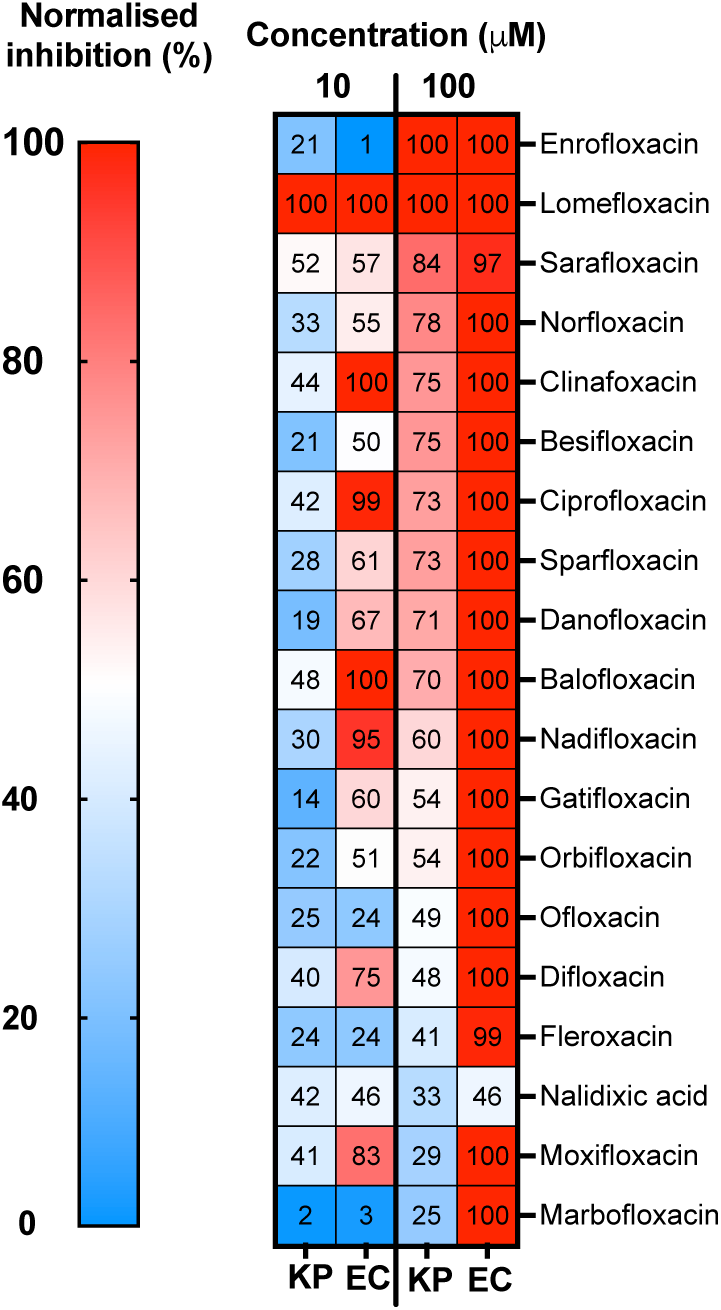
A heatmap of fluoroquinolone and nalidixic acid analogues inhibiting the *E. coli* and *K. pneumoniae* CFE system. Two concentrations at 10 μM and 100 μM were spiked into the reactions. Data is represented by normalised inhibition (%) of the CFE reaction (*n* = 2 independent experiments). Abbreviations: *K. pneumoniae* ATCC 13882 (KP) and *E. coli* MG1655 (EC)

### Membrane-permeabilised cells show similar antimicrobial inhibition to CFE assays

To quantify antibiotic activity between CFE and cell-based growth-inhibition assays, we selected 15 antibiotics based on differences in potency in the CFE assays and intrinsic target (**Figure 5-6**). We selected ten compounds from hits identified from the primary assay that were above the Sigma threshold. These were azithromycin, retapamulin, valnemulin, thiostrepton, kanamycin, rifampicin, spectinomycin, paromomycin, erythromycin and tetracycline. We also included four known antibiotics that were below the Sigma threshold in the CFE assay. They were tigecycline, amikacin, chloramphenicol, and clindamycin, which all target the ribosome. Additionally, bacitracin zinc was added as a non-specific CFE inhibitor since it targets cell wall biosynthesis and was active in CFE at high concentrations. Additionally, to explore limitations in antibiotic transport, we included polymyxin nonapeptide (PMBN) in whole cell MIC assays to increase antibiotic import into the cells. We then conducted full IC and MIC bioactivity assays and quantified the IC_50_/IC_90_ and MIC_50_/MIC_90_ values (**Figure 6a**) for these 15 antibiotics using a 11-point serial dilution from 100 μM in 2% (v/v) DMSO. Here the MIC_50_/MIC_90_ refer to the concentration of compound needed for 50%/90% growth inhibition based on the sigmoidal curve. This was done for comparison between the two assays. In addition, we also generated whole cell bioassay (MIC) data for the laboratory strains *E. coli* MG1655 and *K. pneumoniae* ATCC 13882, and the clinical isolates, *K. pneumoniae* M6, *K. pneumoniae* NCTC 13368, *A. baumannii* ATCC 17978, *E. coli* NCTC 12923, and *A. baumannii* AYE as a relevant comparison between a range of Gram-negative bacteria and clinical isolates. The data for these extended strains is shown in **Table S3**, which reveals a general trend in antibiotic resistance for all the clinical isolates in comparison to *E. coli* MG1655 and the *K. pneumoniae* ATCC 13882 strain.

**Figure 5.**
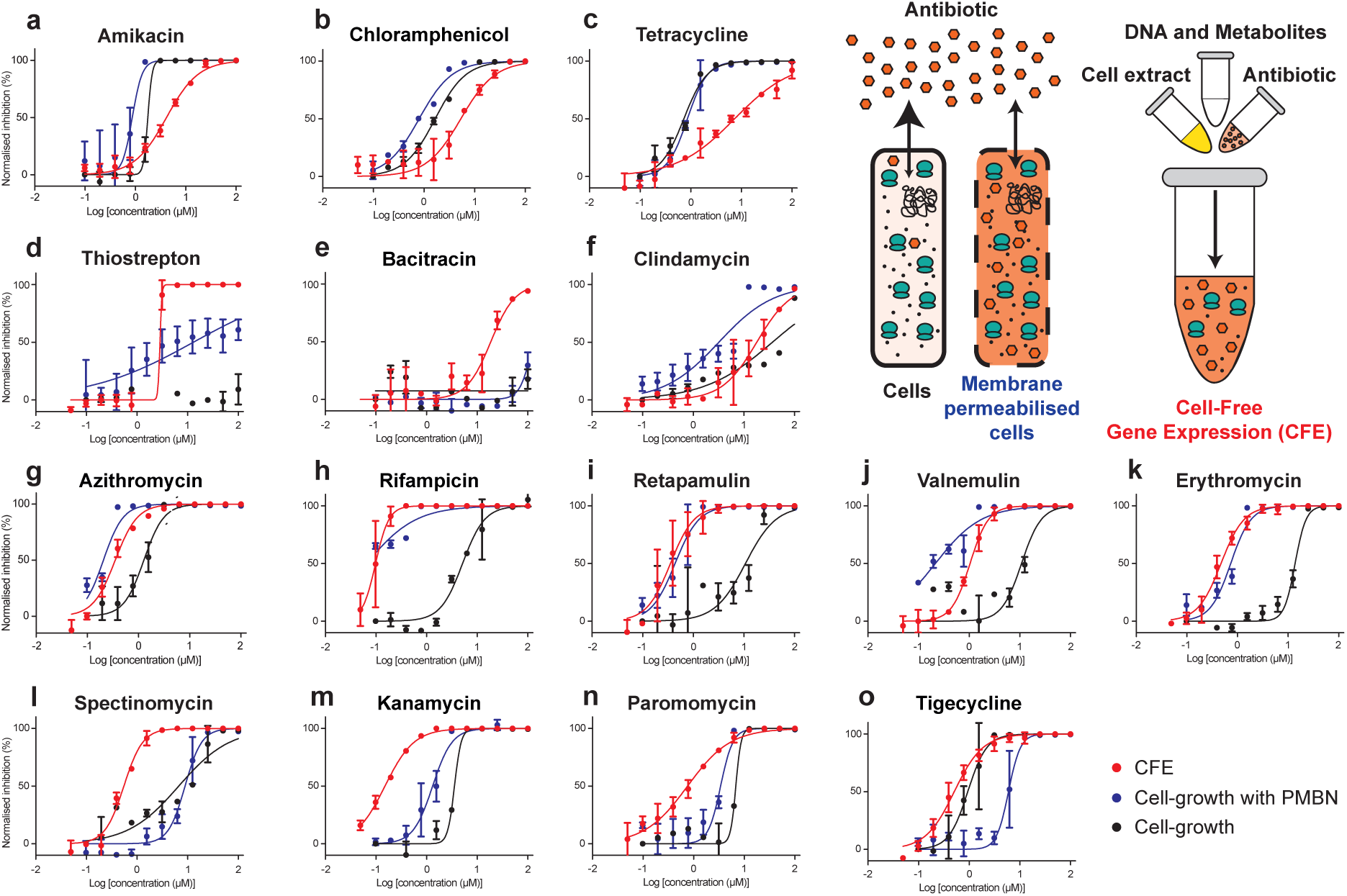
Antibiotic inhibition of *K. pneumoniae* cell-free and whole cells – part I. Antibiotics displaying a similar activity profile between the assays were grouped as follows: (**a-c**) Compounds that were more sensitive in whole cells; (**d-f**) Compounds with weak activity in cells; (**g-k**) Compounds with similar activity in cell-free and cells treated with PMBN; (**i-o**) Compounds with at least an order of magnitude higher sensitivity in cell-free. The data is normalised to provide a relative comparison of Log(concentration) versus normalised inhibition (%) between whole cell and cell-free assays. Data is presented as an average and error bars represent standard deviation. CFE data is the mean and standard deviation of two independent experiments, each with three technical repeats. Each whole cell assay data is mean and standard deviation from two independent experiments).

**Figure 6.**
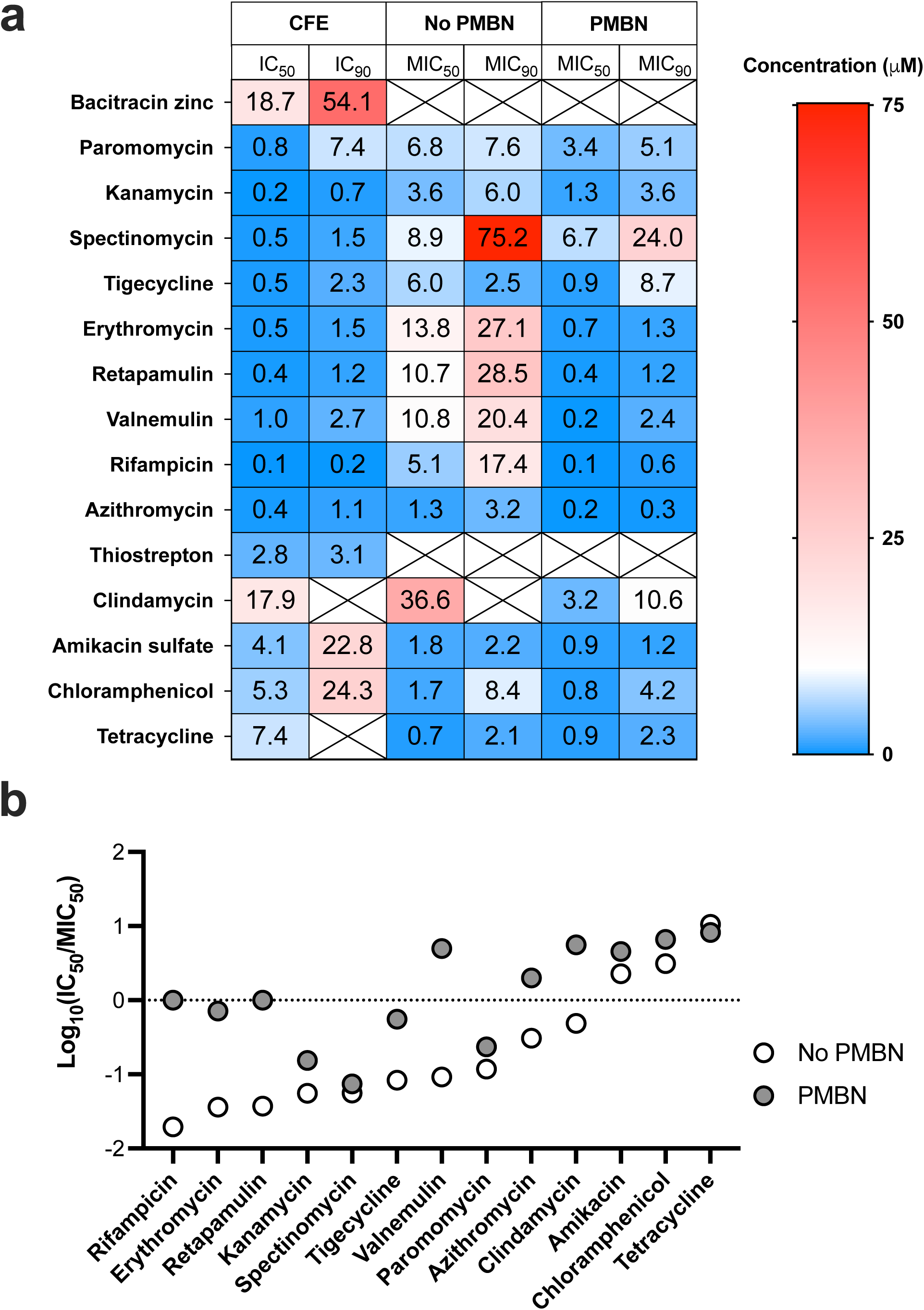
Antibiotic inhibition of *K. pneumoniae* cell-free and whole cells – part II. **(a)** Summary of IC_50_ and MIC_50_ values, displayed from Figure 6. Whole cell data was collected in the presence or absence of PMBN. Boxes with crosses represent undetermined inhibition values. (**b**) Rank ordered comparison of Log_10_(IC_50_/MIC_50_) ratios, where the MIC value is compared between the absence (white circle) and presence (grey filled circle) of PMBN. Thiostrepton, clindamycin and bacitracin are removed because they were inactive in the whole cell assays.

Focusing on both the cell and CFE data for *K. pneumoniae* (**Figure 5**), 12 out of 15 compounds displayed a low IC_50_:MIC_50_ ratio (represented as Log_10_ values in **Figure 6b**), with IC_50_ (most values were between 0.2-1 μM) much lower than the corresponding MIC_50_ values (**Figure 5 and 6**). In contrast, three compounds (chloramphenicol, amikacin sulphate and tetracycline) displayed a high IC_50_:MIC_50_ ratio (**Figure 5a-c**). While there are few studies on antibiotic transport, especially for *K. pneumoniae*, our combined data may indicate increased uptake of these compounds. In addition, *E. coli* is known to import tetracycline and chloramphenicol faster than other common antibiotics^2^, with the latter through an active transport system^47^. In contrast, thiostrepton, bacitracin zinc and clindamycin were active in the CFE system but not in cells.

To probe limitations in antibiotic uptake and its impact on the IC_50_:MIC_50_ ratio, we added PMBN to permeabilise the membrane. This sensitised the cells for thiostrepton (1.1 μM MIC_50_) and clindamycin (7.5 μM MIC_50_), but not bacitracin zinc (**Figure 5d-f**). These data suggest potential transport limitations for these antibiotics. Interestingly, despite inactivity in the cells, bacitracin zinc had a modest IC_50_ value of 18.7 μM. Finally, for azithromycin, rifampicin, retapamulin, valnemulin, erythromycin, spectinomycin, kanamycin, paromomycin and tigecycline, the IC_50_ value was at least an order of magnitude lower than MIC_50_ (**Figure 5g-o**). For seven of these compounds, the MIC_50_ value decreased upon addition of PMBN (**Figure 6**). For five compounds within this set, the dose-response curve closely overlaps for the CFE and PMBN data (**Figure 5g-k**). This provides evidence that for these classes of antibiotics, CFE provides a close bioactivity model for the membrane-permeability assay. Put together, the CFE and cell data highlight where there are potential transport limitations for several antibiotic classes (**Figure 6**), with the CFE approach providing a strain specific tool for exploring this fundamentally challenging question.

### Cell-free profiling of predicted AMR genotypes

We hypothesised that CFE offers unique potential to study target mutations associated with AMR in a macromolecular environment that mimics the cell, without the competing influence of the membrane influx/efflux barriers. However, due to clear limitations of this approach, our initial attempts to do this with CFE systems derived from the clinical isolates ST258-T1b and NJST258-1 was not successful (**Figure S14**). To overcome the issues that we initially encountered, it was desirable to create stable isogenic strains to enable us to compare the effect of resistance-causing mutations in the same strain background. Therefore, we applied adaptive laboratory evolution to generate stable mutants^48^. We serially passaged the ATCC 13882 strain against a selection of antibiotics that were characterised in the CFE and whole-cell assays (**Table S3**). ATCC 13882 was serially passaged with increasing concentrations of antibiotics, starting at 0.25xMIC and doubling every 2 days, until the bacteria were exposed to 4xMIC. Then, the strains were passaged 10x in the absence of antibiotic and the elevated MICs were retained in each case. We then analysed each strain by next-generation sequencing (NGS) and whole genome sequencing (WGS) analysis.

As expected, most antibiotic-resistant strains (e.g., tetracycline, kanamycin) accumulated multiple mutations, some in transport proteins (**Table S3**). However, the rifampicin, chloramphenicol and valnemulin resistant strains contained only one or two genomic mutations, which we prioritised for further study. Focusing on these strains for comparability to the ATCC 13882 strain, the MIC increased from 25 μM for valnemulin and 12.5 μM for rifampicin and chloramphenicol to >200 μM for all three antibiotics. These adapted strains are referred to as resistant to rifampicin (Rif^R^), valnemulin (Val^R^) and chloramphenicol (Chl^R^) derivatives of ATCC 13882. From WGS analysis, the Rif^R^ strain carries a variant (H526L) in RNA polymerase beta subunit (*rpoB*), a mutation also observed in *Mycobacterium tuberculosis* Rif^R^ clinical isolates^49, 50^. H526L is predicted to reduce the binding affinity of rifampicin to RpoB^49^. The Val^R^ strain had a single point mutation (AAC>TAC) at position 445 for *rplC*, which encodes the 50S ribosomal protein L3. Variants at the N149 position are associated with increased valnemulin and linezolid resistance in *E. coli*, with the L3 subunit located close to the peptidyl transferase centre (PTC) in the 50S large ribosomal subunit and the linezolid/valnemulin binding site^51, 52^. Finally, the Chl^R^ strain carries a 24-bp repeat (duplication of amino acids 62-69) in an efflux pump regulator (OqxR) and a I264L variant encoded by a mutated (A790C) hypothetical gene [Locus_tag: WM93_12915, annotated as a histone-like deacetylases (HDAC), class II in related *K. pneumoniae* strain ATCC 35657].

To test the antibiotic-resistant mutant strains for CFE activity and for relative changes in protein synthesis in response to the corresponding antibiotic, cell extracts were isolated (two biological repeats) and an IC_50_ was determined for each specific antibiotic and compared to the ATCC 13882 CFE system. Additionally, we also monitored mScarlet-I synthesis in real-time, in the absence of antibiotics (**Figure 7a**). First, the Rif^R^ variant (RpoB H526L) displayed a 58-fold increase in IC_50_ (*p* value = 0.0201) for rifampicin, compared to the parental strain (**Figure 7b-c**). There were no apparent changes in protein synthesis rates between the ATCC 13882 and Rif^R^ CFE system, suggesting RpoB H526L does not alter the rate of RNAP significantly. This was consistent across two biological repeats, suggesting that the H526L variant directly confers resistance to rifampicin, which agrees with previous studies^49, 50^. For the Val^R^ CFE system, surprisingly there was no apparent change in IC_50_ when valnemulin was added (**Figure S18)**. While there was a slight delay in protein synthesis in comparison to the ATCC 13882 CFE system, the final yield of protein was similar (**Figure S19**). The unaltered IC_50_ value was unexpected, given that the RplC N149Y variant occurs close to the valnemulin and linezolid binding site, and that this was the only mutation identified from NGS analysis of this strain. Last, the Chl^R^ CFE system showed a significant increase in the IC_50_ value for chloramphenicol determined at 16.3 μM (*p* value = 0.0154), in comparison to 5.3 μM measured for the ATCC 13882 strain (**Figure 7d-e**). This might implicate that the histone-like deacetylase (HDAC) homologue (WM93_12915) somehow exerts a modest effect on chloramphenicol inhibition in the CFE system, through an unknown mechanism, while the greater increase in the MIC value is likely due to the upregulation of efflux pumps through the mutation in the *oqxR* regulator. Finally, we also tested the ATCC 13882, Rif^R^ and Chl^R^ strains for potential changes in virulence using the non-animal infection model, *Galleria mellonella*^5^ (**Figure 7f-g**). This insect model provides a relatively simple *in vivo* system to perform virulence tests. To do this, we infected *G. mellonella* with a ‘high’ or ‘low’ inoculum (see methods) of the *K. pneumoniae* strains and recorded survival over time. The Rif^R^ mutant displayed a significant loss in virulence at both inoculums compared to the parent strain (*p* value <0.0001), while the Chl^R^ mutant shows increased virulence, which is defined as significant at the higher inoculum (*p* value <0.05). Together with the rest of the findings, this virulence assay provides added insight into the potential impact of these AMR variants discussed, from an *in vivo* perspective.

**Figure 7.**
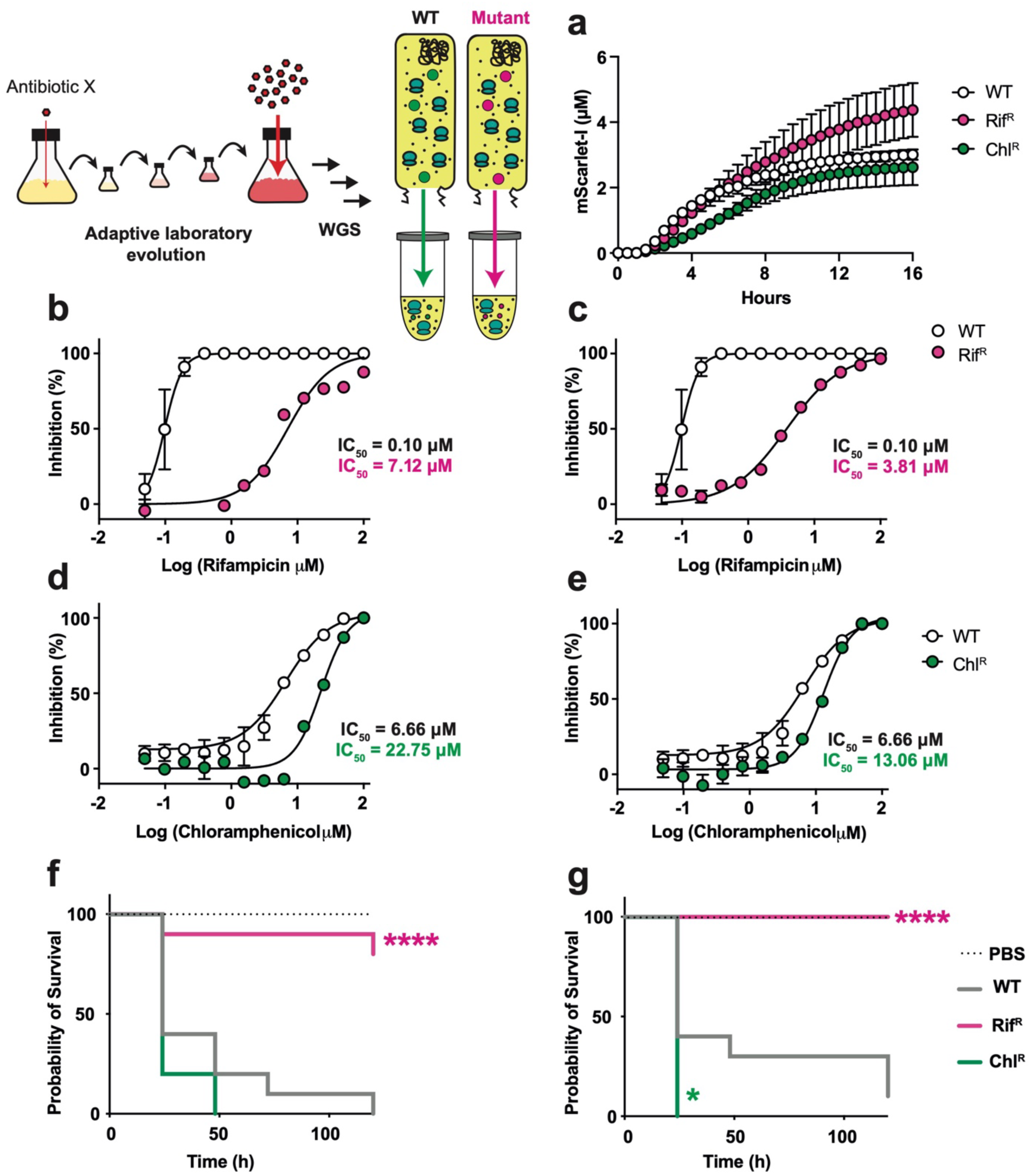
A laboratory adaptation, evolution, and cell-free characterisation platform for quantifying the effects of antibiotic resistance at the level of *K. pneumoniae* CFE. (**a**) Real-time protein synthesis of mScarlet for the wild type (WT), Rif^R^ and Chl^R^ mutants. (**b-c**) IC_50_ data of two biological repeats for the Rif^R^ mutant plotted separately versus a representative dataset for the WT in response to rifampicin dosing. CFE data for WT is mean and standard deviation of two independent experiments(**d-e**) IC_50_ data of two biological repeats for the Chl^R^ mutant plotted separately versus a representative dataset for the WT in response to chloramphenicol dosing. CFE for WT is mean and standard deviation of two independent experiments. (**f-g**) Killing assay of *G. mellonella* infected with *K. pneumoniae* WT, Rif^R^ and Chl^R^ mutants with either a low (f panel) or high (g panel) infective dose (see methods).

## Discussion

AMR is a global pandemic projected to cause 10 million annual deaths by 2050, with a total economic cost of $100 trillion^53^. Therefore, it is vital to find new ways to explore AMR, as well as develop novel counter strategies (e.g., non-standard antimicrobials, phage therapy, host-directed therapeutics) to fight resistance. By removing the cell wall and membrane, we establish a *K. pneumoniae* CFE system, which provides a fast and safe model to study AMR genotypes in a WHO priority pathogen. While a similar approach can be performed with *in vitro* approaches (e.g., purified ribosomes, enzyme complexes and mutants), target inhibition of an isolated protein has so far failed to deliver new antibiotic candidates, despite extensive HTS studies^54^. Alternatively, we suggest CFE systems are more straightforward, and provide the native macromolecular proteins and metabolites unique to the original host cell. This also provides a safe test tube mimic of a living cell.

Antibiotics were discovered by industrial HTS of natural product and synthetic libraries during the 1940-60s^53, 55^. Due to a range of factors, including AMR and economics, this discovery trail has long since faded^56, 57^. Recently, there has been new investment in antimicrobial development around small-medium enterprises^58^. Towards this need, CFE systems offer a low cost model with high sensitivity for identifying antimicrobial compounds from natural product and synthetic sources^15, 17^. However, with the exception of a bespoke *S. pneumoniae* CFE system^17^, early IVT systems generally use established models such as *E. coli* and *B. subtilis*^13, 14^. Recent advances in cell-free systems provide new opportunities to study almost any cell type that can be grown in the laboratory including from prokaryotes^19–24^ and eukaryotes^59–61^. After the initiation of our study, a *K. pneumoniae* CFE system was recently released^62^, but alternatively this study focused on the potential of this cell-free system for biotechnology and protein production. In contrast, we provide a *K. pneumoniae* CFE system for studying antimicrobials and AMR factors intrinsic to this microbe. Importantly, while developing this screening platform, we apply recommended HTS standards to validate our study^43^. Key features of our CFE system include strong performance for the S:B ratio and robust Z’-factor score. We estimate that for HTS screening, our CFE platform costs less than $0.01 per drug. For comparison, most HTS campaigns cost $0.10-0.50 for single target enzyme assays, or $5-10 for whole cell targets^63^. Moreover, our *K. pneumoniae* CFE system also showed remarkably high sensitivity and specificity, with minimal non-specific hits in comparison to a *E. coli* CFE benchmark. This also identified novel features of the assay, such as its ability to identify inhibitors of indirect targets (e.g., DNA gyrase/topoisomerase).

By extending our studies to whole cell MIC assays with a wide range of clinical antibiotics, we highlight the potential benefits for quantifying the on target inhibitory effects of antibiotics that are compromised in whole cell assays by issues due to antibiotic transport. Antibiotic potency is limited by flux across the cell envelope and overall concentration inside the cell. Understanding antibiotic transport remains a significant barrier in AMR research, with studies limited to established models (i.e., *E. coli*). It is especially challenging to quantify antibiotic influx/efflux for Gram-negative bacteria because of two membrane layers and the periplasmic space^2^. Our data shows 12 out of 15 antibiotics display a low IC_50_:MIC_50_ ratio between the CFE and whole MIC assays for *K. pneumoniae*. As an exception, chloramphenicol and tetracycline display a high IC_50_:MIC_50_ ratio, while bacitracin (cell-wall biosynthesis inhibitor) was inactive in whole MIC assays but displayed non-specific inhibition in the CFE system. Chloramphenicol and tetracycline are known to have a high accumulation rate for *E. coli* MG1655 compared to other common types of antibiotics^2^, which for chloramphenicol, is linked to active uptake mechanisms^47^. Unfortunately, there are no equivalent studies on antibiotic transport in *K. pneumoniae* to compare to. Instead, we suggest our antimicrobial measurements (and IC_50_:MIC_50_ ratios) may improve the understanding of the balance between target inhibition, efflux liability and membrane permeability effects observed between the different assays.

To the best of our knowledge, we uniquely provide a CFE system which allows the study of AMR from an intracellular perspective. While the cell envelope plays a major role in AMR, we provide a system to directly study how resistance mutations, associated with intracellular components, alter gene expression. A CFE system provides a core proteome catalysing coupled transcription-translation, energy regeneration and amino acid flux. There are also potentially other essential processes, such as RNA modification, protein degradation, folding chaperones, that remain to be studied within the context of a cell-free extract. Herein, we study three laboratory-adapted *K. pneumoniae* strains with either single or double genomic mutations associated with AMR. While the antibiotics used are not clinically relevant for *K. pneumoniae* treatment, they were selected based on their targets (e.g., ribosomes, RNAP) and low MIC value observed in the whole cell characterisation. From this we provide proof of principal data for how individual mutations influence gene expression and AMR. We confirm previous findings from Rif^R^ *M. tuberculosis*^49, 50^ that the RNA polymerase beta subunit H526L variant confers strong CFE resistance to rifampicin. Interestingly, this variant also showed reduced virulence in a *in vivo* wax moth model suggesting this mutation compromises pathogenicity. For the Chl^R^ strain, it is known that HDAC have been implicated in the establishment of persistence and antibiotic tolerance in *Burkholderia thailandensis*^64^, which is consistent with the CFE phenotype observed here. If CFE can be used to study this complex regulation of bacterial metabolism, then this might be a valuable tool for developing new adjunct therapies to prevent antibiotic tolerance. Since the HDAC is a hypothetical protein, the precise intracellular resistance mechanism remains unclear at this stage. In contrast, the >16-fold increase in MIC appears to be linked to the 24-bp repeat (duplication of amino acids 62-69) within the gene encoding the OqxR regulator a protein already linked to significantly up-regulated expression of the OqxAB efflux pump^65^. Finally, for a Val^R^ strain, there was no apparent change in IC_50_ value in response to valnemulin. This was an unexpected since the mutation site, L3 N149Y, is located close to the PTC, close to the valnemulin/linezolid binding region. While this specific mechanism remains unknown, overall, our findings demonstrate that CFE provides a powerful approach to investigate the effects of specific mutations in known antibiotic targets and may clarify specific functions which might not be evident from either whole cell or single target studies. Since the system provides the transcription-translation steps and metabolic processes, unique to *K. pneumoniae*, the effects of mutations associated with these processes can be monitored with the final reaction – i.e., protein synthesis. We highlight the RpoB H526L variant specifically, since this provides a clear phenotype, which is also consistent across biological repeats. As such, we provide a significant new tool to enable the study of AMR and to assist in identifying new antimicrobial agents and resistance mechanisms.

Overall, our CFE model provides a highly selective and sensitive evaluation of specific antimicrobial targets and genetic determinants associated with AMR in *K. pneumoniae*, a major Gram-negative bacterial pathogen. While initial preparation in a CL2 laboratory is required, after processing, the extracts are stable and non-living, and thus safe to study in a CL1 lab. This added dexterity which CFE provides, leads to a significant new research tool for AMR research, which together with recent advances in cell-free systems, offers new opportunities to explore infectious diseases.

## Acknowledgements

SJM and CMS acknowledge Global Challenges Doctoral Centre (GCDC) at the University of Kent for PhD studentship funding for KC. SJM and CMS would like acknowledge Dr Gary Robinson (University of Kent) for access to the Selleck FDA library. Work at UKHSA was supported by Grant in Aid for Open Innovation in AMR (Project 111742). J.A.B. laboratory is supported by grants from the Medical Research Council (MR/V032496/1) and Biotechnology and Biological Sciences Research Council (BB/V007939/1).

## Author contributions

KC, CMS, CH, JMS and SJM designed the study; KC, CH, MW, LN, and TH performed the experiments; KC, CH and MW analysed the data; AT, RME and JAB contributed resources and consulted on the study; KC, CH, CMS, JMS and SJM wrote the paper.

## Methods

### Molecular biology and bacterial strains

*E. coli* MG1655 and DH10β (NEB) strains and all molecular biology was performed as previously described^38^. *K. pneumoniae* strain ATCC 13882 was purchased from Deutsche Sammlung Von Mikroorganismen and Zellkulturen (DSMZ). *K. pneumoniae* clinical strains ST258-T1b and NJST258-1 were previously described^66, 67^. The strains were maintained and grown in standard Luria-Bertani, or specific media described within the supporting information. Clinical strains used for minimum inhibitory concentration assays (*K. pneumoniae* M6 and NCTC 13368, *A. baumannii* ATCC 17978 and AYE (BAA-1710), *E. coli* NCTC 12923) were acquired from ATCC or NCTC and maintained on Tryptic Soy Broth. All mutant strains of *K. pneumoniae* ATCC 13882 were generated at UKHSA (**Table S3**) using laboratory adapted evolution and sequenced. Further details are provided within the methods.

### Growth measurements

A single colony of *K. pneumoniae* ATCC 13882 was inoculated into 5 mL of desired media and incubated at 30°C, 200 rpm for 16-18 h. The pre-culture was diluted 1:100 in 50 mL of fresh sterile media in a 250 mL conical flask. Optical density at 600 nm (OD_600_) was recorded in a UV-Vis spectrophotometer (Agilent, Cary 60), using sterile media as blank. Subsequently, OD_600_ measurements were taken every h (the culture was diluted 1 in 10 when OD_600_ ≥ 1) and a growth curve was plotted and visualised using GraphPad Prism version 9.0.

### *K. pneumoniae* cell extract preparation

Cell extract preparation consisted of the following steps: cell growth and harvest, cell washes with S30A buffer, lysis by sonication, run-off reaction, and, when specified, dialysis. The protocol for crude extract preparation was adapted and modified from previous studies^17, 37^. *K. pneumoniae* ATCC 13882 was grown overnight on LB agar at 30°C. A single colony was then inoculated into 5 mL of LB and grown at 30°C, 200 rpm for 6-8 h (unless specified otherwise). 50 mL of fresh sterile media was then inoculated with 250 μL of pre-culture and incubated at 30°C, 200 rpm for 14-16 h. The following day, the overnight culture was diluted 1:100 in 500 µL growth media in Ultra-Yield™ (Thomson, USA) baffled flasks and incubated at 30°C, 220 rpm until the OD_600_ reached 2.5-3.0. Cells were transferred into pre-chilled 250 mL centrifuge bottles and centrifuged (10,000 x *g*, 4°C, 12 min). The supernatant was discarded. Then, cells were washed with 20 mL S30A buffer, recombined into a 30 mL centrifuge bottle and centrifuged (15,000 x *g*, 4°C, 12 min). The supernatant was carefully discarded, and wash step was repeated. The cell pellet was resuspended, aliquoted into 2 mL polypropylene (PP) tubes and centrifuged (18,000 x *g*, 4°C, 12 min). The residual supernatant was discarded using a pipette tip. Using a pipette with the tip cut to increase bore size, the cell pellets were combined into a 50 mL PP tube for cell lysis by sonication. Cells were sonicated in an ice-water bath, in either a 50 mL PP tube or aliquoted into 1.5 mL PP tubes. Settings used were: 20 kHz frequency, 50% amplitude, 10 sec pulse on time, 10 sec pulses off time, 500 J energy input/mL of wet-cell pellet. The lysate was centrifuged (18,000 x *g*, 4°C, 10 min), the supernatants were pooled and then aliquoted into 1.5 mL PP tubes. The lysates were incubated at 37°C for 60 min (unless otherwise stated) and then clarified by centrifugation (18,000 x *g*, 4°C, 10 min). When required, clarified lysate was pooled and dialysed for 3 hr at 4°C in S30B buffer. The extract was subsequently clarified by centrifugation (18,000 x *g*, 4°C, 10 min), aliquoted into 200-500 μL fractions and stored at -80°C. Total protein concentration of the crude extract was estimated using Bradford assay with a BSA standard, as previously described^39^. Using this method, the typical protein concentration of crude extracts from *K. pneumoniae* was 20-24 mg/mL.

### K. pneumoniae CFE reactions

The CFE reactions consisted of the following final concentrations (unless otherwise stated): 50 mM HEPES, 30 mM 3-PGA, 1.5 mM ATP, 1.5 mM GTP, 0.75 mM CTP, 0.75 mM UTP, 1 mM amino acids, 0-10 mM magnesium-glutamate, 0-200 mM potassium-glutamate, 3% (w/v) polyethylene glycol 8000, 0.2 mg/ml *E. coli* tRNA, 0.26 mM CoA, 0.33 mM NAD, 0.75 mM cAMP, 0.068 mM folinic acid, 1 mM spermidine and 10 nM plasmid DNA. All chemicals were prepared, as previously described^37^. For end-point measurement of protein synthesis, reactions were performed in 22 or 33 µL reactions, in PP tubes as specified. Reactions were incubated at 28°C, 250 rpm for 6 h or overnight (14-16 h). For time-course measurement, the reaction mixture was transferred as 10 µL technical replicates in a 384-well black, clear flat-bottom microplate. Fluorescence measurements (excitation filter 544 and emission filter 620 nm, 10 nm bandwidth) were recorded in a microplate reader every 10 min or after end-point incubation at 28°C with 5 s of 250 rpm orbital shaking prior to measurement. Calibration standards of purified N-terminal His_6_-tagged mScarlet-I were prepared in the reaction mixture as described previously^39^. For *E. coli* MG1655 CFE, extracts were prepared as previously described^40^.

### Whole cell assays with minimum inhibitory evaluation (MIC assay)

In this study, the MIC_50_ and MIC_90_ values are defined as the concentration of compound needed to cause 50 and 90% inhibition respectively. A single colony was taken from a freshly grown agar plate of each strain and inoculated into 3 mL of sterile tryptic soy broth. The pre-cultures were incubated at 37°C for 16-18 h with shaking at 200 rpm and then diluted to an OD_600_ of 0.01, equivalent to 1 x 10^6^ colony-forming units (CFU) per mL. This was further sub-cultured into a sterile clear flat-bottom 96-well plate to 5 x 10^5^ CFU per mL and dosed with one compound per plate. Each compound was serially diluted in 100 µL of TSB broth to which 100 µL of culture was added. Wells containing media and compound dilution were included as negative control while wells with bacterial strain only were used as positive control. The plates were incubated at 37°C. During this incubation, bacterial growth was determined by reading absorbance at 600 nm, on a CLARIOstar microplate reader (BMG, LabTech) with an attached plate-stacker enabling measurements of optimal density at 600 nm (OD_600_) every 30 min for a total of 20 h. Background signal was eliminated by subtracting absorbance in blank wells. Data was normalised to the OD_600_ of positive control wells or wells with lowest concentration of compound (whichever was higher) and expressed as percentage growth. MIC cut-off values were determined according to Clinical and Laboratory Standards Institute (CLSI) guidelines. Data is fitted to either a symmetric (4-parameter) or asymmetric (5-parameter) logistic regression model and visualised using GraphPad Prism version 9.0.

### CFE screening

The Selleck FDA-approved drug screening library containing 1175 compounds (10 mM in DMSO) was screened. The drugs were tested at an initial concentration of 10 μM with 1% (v/v) DMSO in a *K. pneumoniae* ATCC 13882 CFE reaction, at a total volume of 22 μL in 0.2 mL PCR tubes. Samples were mixed by briefly vortexing for 5 s. Two 10 μL aliquots were transferred to a 384-well microtiter plate as 10 μL duplicate measurements. Plates were sealed using a gas-permeable optically clear adhesive film. They were briefly centrifuged (200 × *g*, 10 s). Reactions were run for 16 h at 30°C. To validate the performance of the CFE reaction, a robust Z’ (RZ) factor analysis was performed, as previously described^23^. Controls included negative (no DNA), positive (DNA only), 1% (v/v) DMSO (DNA with DMSO) and 10 µM rifampicin (with DNA). A secondary screen was performed by testing potential hits (with an initial % inhibition between 20 and 90%) at 100 µM in 1% (v/v) DMSO.

### CFE IC_50_ characterisation

Select compounds (see results and supporting data) were tested between a range of 0.04 μM to 100 μM via serial dilution in 1% (v/v) DMSO, to determine half maximal inhibitory concentration (IC_50_) values. CFE reactions were performed as described in the primary screen, in three technical replicates and using two different extracts. Data is fitted to either a symmetric (4-parameter) or asymmetric (5-parameter) logistic regression model and visualised using GraphPad Prism version 9.0.

### Laboratory-adapted evolution

The *K. pneumoniae* ATCC 13882 strain was serially passaged against a range of antibiotics selected from the CFE and whole-cell assays (**Table S3**). *K. pneumoniae* ATCC 13882 was serially passaged with increasing concentrations of antibiotics, starting at 0.25xMIC and doubling every 2 days, until the bacteria were exposed to 4xMIC. Then, the strains were passaged 10x in the absence of antibiotic and the MICs were measured again and the elevated MICs were retained in each case.

### WGS of *K. pneumoniae* strains

Genomic DNA was purified using a Wizard genomic DNA purification kit (Promega). DNA was tagged and multiplexed with the Nextera XT DNA kit (Illumina). Whole-genome sequencing of *K. pneumoniae* isolates was performed by UKHSA-GSDU (UK health Security Agency Genomic Services and Development Unit) on an Illumina (HiSeq 2500) with paired-end read lengths of 150 bp. A minimum 150 Mb of Q30 quality data were obtained for each isolate. FastQ files were quality trimmed using Trimmomatic. SPAdes 3.15.3 was used to produce draft chromosomal assemblies, and contigs of less than 1 kb were filtered out. FastQ reads from antibiotic-exposed isolates were subsequently mapped to the genome sequence of pre-exposed strain ATCC 13882 using BWA-MEM (version 0.7.17). Bam format files were generated using Samtools (version 1.6), VCF files were constructed using SNIPPY (Version 4.6.0) GATK2 Unified Genotyper (version 0.0.7) (50). They were further filtered using the following filtering criteria to identify high-confidence SNPs: mapping quality, _60; genotype quality, 40; variant ratio, _0.9; read depth, _10. All the above-described sequencing analyses were performed using Galaxy (www.useglaxy.org). BAM files were visualized in Integrative Genomics Viewer (IGV) version 2.12.2 (Broad Institute).

### *G. mellonella* killing assays

Wax moth larvae (*G. mellonella*) were purchased from Livefood UK and were maintained on wood chips in the dark at room temperature. The larvae were stored for no longer than 2 weeks. Bacterial infection of *G. mellonella* was carried out essentially as described^5^. K. pneumoniae were diluted in PBS to an OD_600_ of 0.002 or 0.01, representing low or high doses (in reference to **Figure 7**). Determination of intracellular bacterial numbers was performed as previously described^68^, except that appropriate dilutions were plated out onto LB agar plates supplemented with the selection antibiotic, which were incubated overnight at 37 °C to allow the bacteria to grow. Data analysis and visualisation was performed using GraphPad Prism version 9.0.

