## Supporting Information for "A cell-free strategy for profiling intracellular antibiotic sensitivity and resistance"

**Supporting Information – Table of Contents**


| | |
|---|---|
| Figure S1 | Growth optimisation I |
| Figure S2 | Growth optimisation II |
| Figure S3 | Growth optimisation III |
| Figure S4 | Time course showing synthesis of the mScarlet-I protein in *K. pneumoniae* ATCC 13882 CFE from extracts harvested at different growth stages |
| Figure S5 | Effect of growth temperature on *K. pneumoniae* ATCC 13882 CFE activity |
| Figure S6 | Optimisation of sonication energy input for *K. pneumoniae* ATCC 13882 CFE activity |
| Figure S7 | Optimisation of incubation (“run-off reaction”) post cell-lysis for *K. pneumoniae* ATCC 13882 CFE activity |
| Figure S8 | Effect of dialysis on *K. pneumoniae* ATCC 13882 CFE activity |
| Figure S9 | Optimisation of DNA concentration for optimal *K. pneumoniae* ATCC 13882 CFE activity |
| Figure S10 | Comparing the effect of nucleotide triphosphates (NTPs) or monophosphates (NMPs) as primary energy source in *K. pneumoniae* ATCC 13882 CFE activity of extracts harvested at different growth stages |
| Figure S11 | Optimisation of Mg-glutamate and K-glutamate concentrations for the *K. pneumoniae* ATCC 13882 cell extract used for library screening experiments |
| Figure S12 | Relative activity (mScarlet-I fluorescence) of extracts before and after filter-sterilisation |
| Figure S13 | Recovery of viable colony forming units from *K. pneumoniae* ATCC 13882 cell-extracts |
| Figure S14 | Antibiotic-resistance profiling for *K. pneumoniae* NJST258-1 |
| Figure S15 | Preliminary antimicrobial inhibition of *K. pneumoniae* ATCC 13882 CFE |
| Figure S16 | Effect of incubation time during reaction setup on assay stability |
| Figure S17 | Effect of DMSO (v/v) concentration on *K. pneumoniae* ATCC 13882 CFE activity |
| Figure S18 | Dose-response curve of valnemulin inhibition of the ATCC 13882 (WT) and Val$^{R}$ CFE systems |
| Figure S19 | Real-time protein synthesis of the ATCC 13882 (WT) and Val$^{R}$ CFE systems |
| Table S1 | Plasmids |
| Table S2 | Extended MIC data on Gram-negative type and clinical strains |
| Table S3 | Whole genome sequence summary of ATCC 13882 antibiotic resistant strains |
| Supporting File 1 | CFE antibiotic screening data |

### Optimisation of the *K. pneumoniae* ATCC 13822 CFE system

To establish a *K. pneumoniae* ATCC 13882 cell-free system, we followed the previous *E. coli* cell-free protocol with some modifications[39]. Cells were grown at 37°C, and harvested at an $OD_{600}$ of 2.5, followed by cell-lysis by sonication, and then finally, extract processing and clarification. Data for ATCC 13882 growth and CFE reaction optimisation is shown in **Figure S1 to S10**. Approximate protein yields for cell extracts were between 20-22 mg/mL, as determined by Bradford assay. Cell-free reactions were prepared with 10 nM plasmid DNA, or without, and incubated for 6-12 hr at 30°C to monitor for either eGFP or mScarlet-I fluorescence.

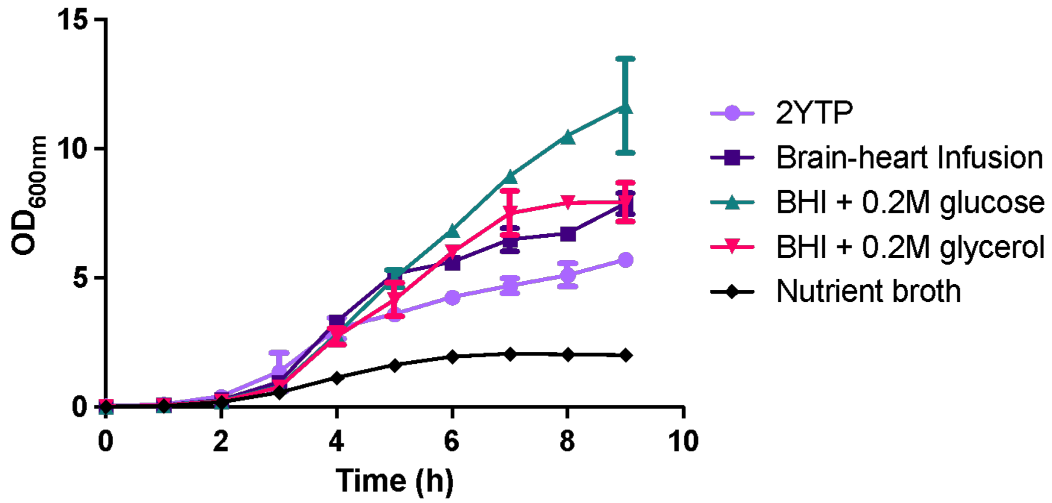


**Figure S1**. Growth optimisation I. Growth curve of *K. pneumoniae* ATCC 13882 in various rich media. Cells were cultivated in 50 mL liquid cultures grown in 250 mL baffled flasks at 30°C, 200 rpm. Data is shown as mean ± standard error of mean (*n* = 2 biological repeats).

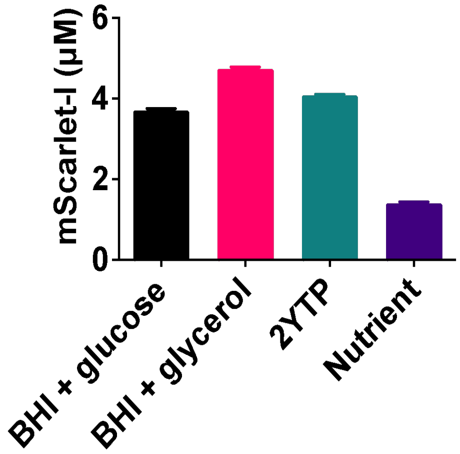


**Figure S2**. Growth optimisation II. *K. pneumoniae* ATCC 13882 CFE activity from extracts generated from different growth media at an $OD_{600}$ of 2.0. Cell-free reaction conditions: 8 mg/mL cell extract, 10 nM pTU1-A-SP44-mScarlet, standard energy solution, incubated at 30°C for 15 hours. Data is shown as mean ± standard deviation of three technical measurements.

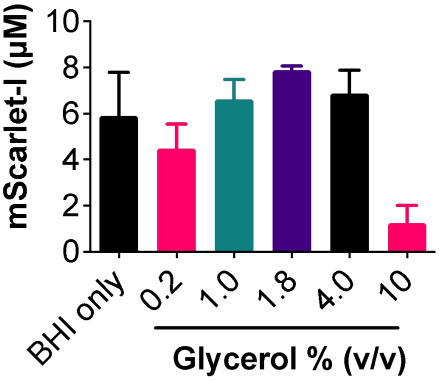


**Figure S3**. Growth optimisation III. *K. pneumoniae* ATCC 13882 CFE activity from extracts grown in BHI supplemented with and without glycerol. Cells were cultivated in 50 mL BHI media with varying concentrations of glycerol at 30°C, 200 rpm, and harvested at an $OD_{600}$ of 2.0. Cell-free reaction conditions: 8 mg/mL cell extract, 10 nM pTU1-A-SP44-mScarlet, standard energy solution and 30°C for 15 hours. Data was shown as mean ± standard error of mean (*n* = 2 biological repeats).

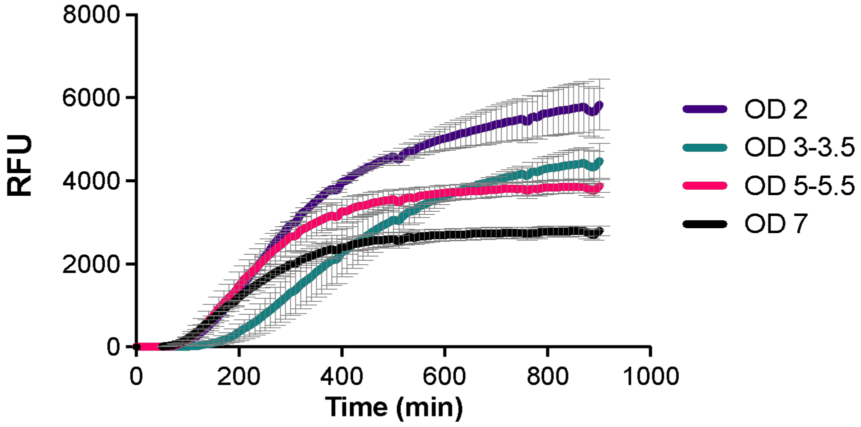


**Figure S4**. Time course showing synthesis of the mScarlet-I protein in *K. pneumoniae* ATCC 13882 CFE from extracts harvested at different growth stages. Cells were grown in 50 mL BHI media with 1.8% glycerol (v/v/) at 30°C, 200 rpm and harvested at the mentioned OD600. Cell-free reaction conditions: 8 mg/mL cell extract, 10 nM pTU1-A-SP44-mScarlet, standard energy solution and 30°C for 15 hours, with fluorescence measurement every 10 min. Data is shown as mean ± standard error of mean (*n* = 2 biological repeats).

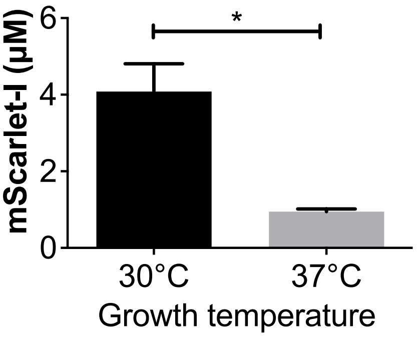


**Figure S5**. Effect of growth temperature on *K. pneumoniae* ATCC 13882 CFE activity. Cells were cultivated from 50 mL cultures in BHI supplemented with 1.8% (v/v) glycerol, at 30 °C or 37 °C until an $OD_{600}$ of 2.0-2.5 was reached. Data is shown as mean ± standard error ($n$ = 2 biological repeats). *$p<0.05$ following a one-tailed unpaired $t$-test.

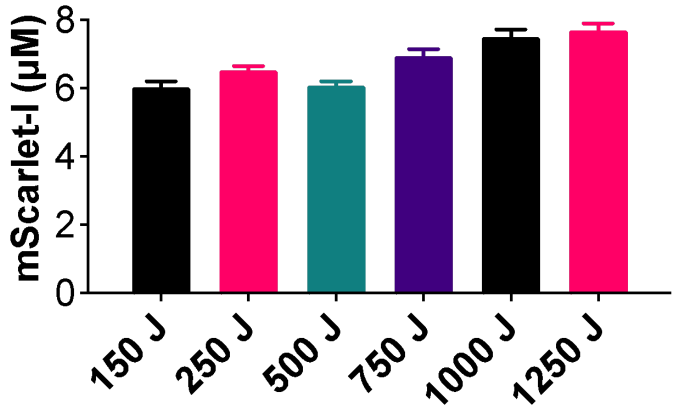


**Figure S6**. Optimisation of sonication energy input for *K. pneumoniae* ATCC 13882 CFE activity. Cell-free reaction conditions: 8 mg/mL cell extract, 10 nM pTU1-A-SP44-mScarlet, standard energy solution and incubated at 30°C for 15 hours. Data is shown as mean ± standard error of mean ($n$ = 2 biological repeats).

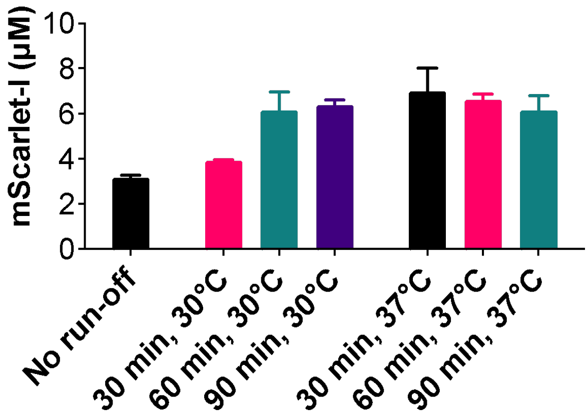


**Figure S7**. Optimisation of incubation ("run-off reaction") post cell-lysis for *K. pneumoniae* ATCC 13882 CFE activity. Cell-free reaction conditions: 8 mg/mL cell extract, 10 nM pTU1-A-SP44-mScarlet, standard energy solution and 30°C for 15 hours. Data is shown as mean ± standard error of mean ($n$ = 2 biological repeats).

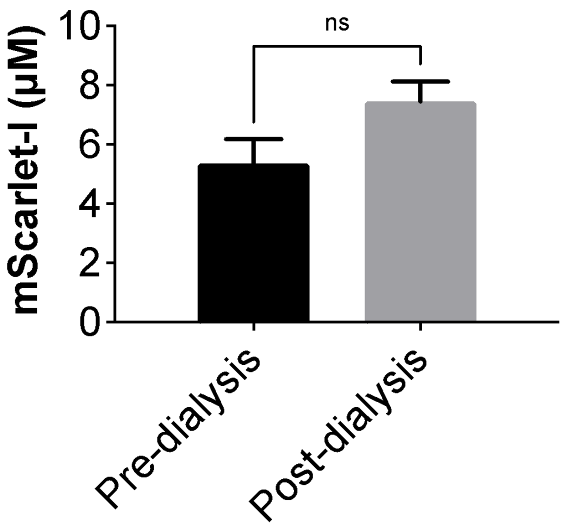


**Figure S8**. Effect of dialysis on *K. pneumoniae* ATCC 13882 CFE activity. Cell-free reaction conditions: 8 mg/mL cell extract, 10 nM pTU1-A-SP44-mScarlet, standard energy solution and 30 °C for 15 hours. Extracts were dialysed against 1 L of S30B buffer for 3 h at 4 °C. Data is shown as mean ± standard error of mean for three biological repeats. P value > 0.05 following a two-tailed paired *t*-test (*n* = 3 biological repeats).

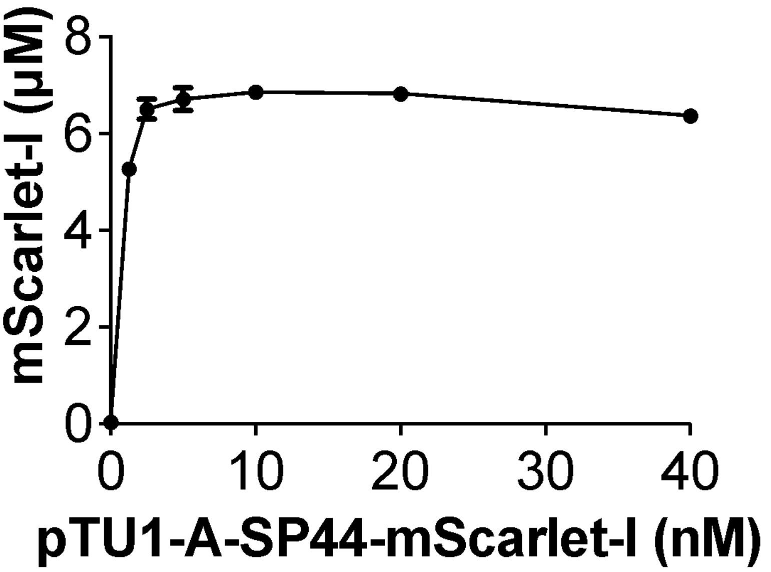


**Figure S9**. Optimisation of DNA concentration for optimal *K. pneumoniae* ATCC 13882 CFE activity. Cell-free reaction conditions: 8 mg/mL cell extract, 0-40 nM pTU1-A-SP44-mScarlet, standard energy solution and 30°C for 15 hours. Data is shown as mean ± standard deviation of one representative dataset with three technical repeats (*n* = 2 biological repeats).

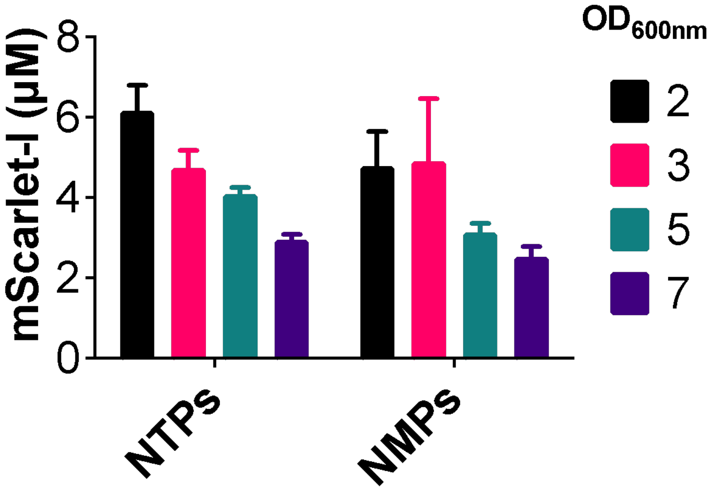


**Figure S10**. Comparing the effect of nucleotide triphosphates (NTPs) or monophosphates (NMPs) as primary energy source in *K. pneumoniae* ATCC 13882 CFE activity of extracts harvested at different growth stages. Cell-free reaction conditions: 8 mg/mL cell extract, 10 nM pTU1-A-SP44-mScarlet, standard energy solution with either NTPs or NMPs and incubated at 30°C for 15 hours. Data is shown as mean ± standard error of mean ($n$ = 2 biological repeats).

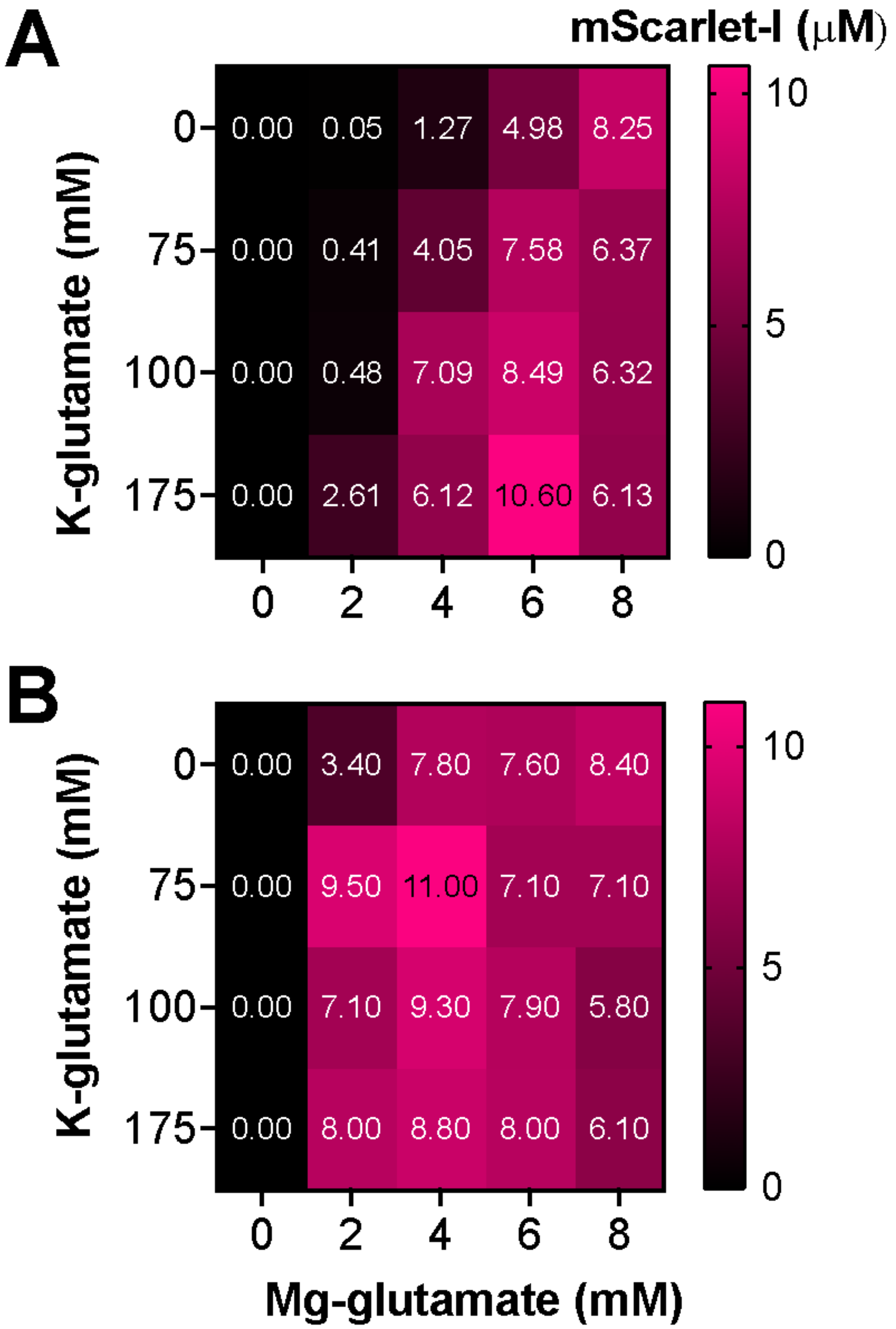


**Figure S11**. Optimisation of Mg-glutamate and K-glutamate concentrations for the *K. pneumoniae* ATCC 13882 cell extract used for library screening experiments. Top (A) and bottom (B) panels show two independent batches of cell extract. Each time a new batch was prepared, this optimisation step was performed to ensure maximal activity and consistency between the experiments. Cell-free reaction conditions: 8 mg/mL cell extract, 10 nM pTU1-A-SP44-mScarlet, standard energy solution and 30°C for 15 hours. Data is shown as mean of three technical repeats.

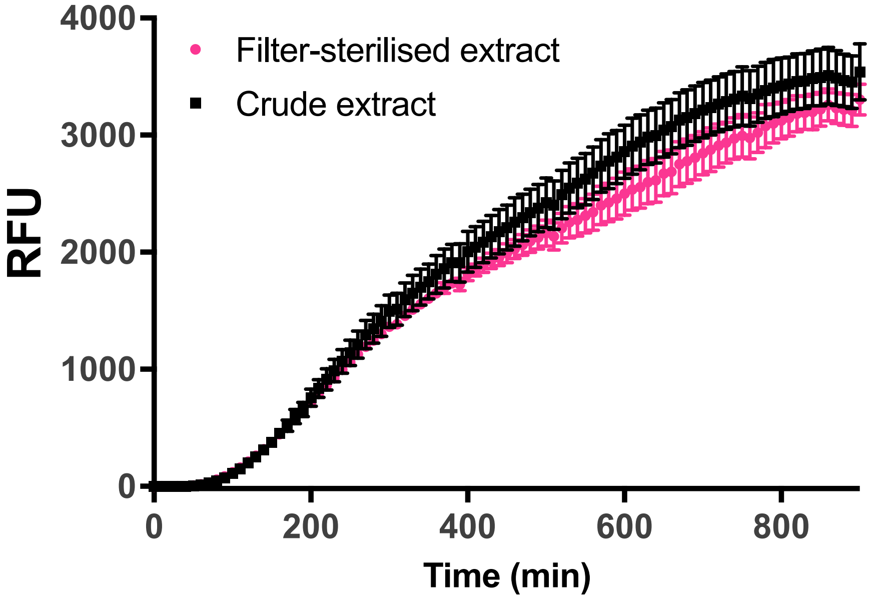


**Figure S12.** Relative activity (mScarlet-I fluorescence) of extracts before and after filter-sterilisation. Reactions were prepared and measured using the standard methods outlined.

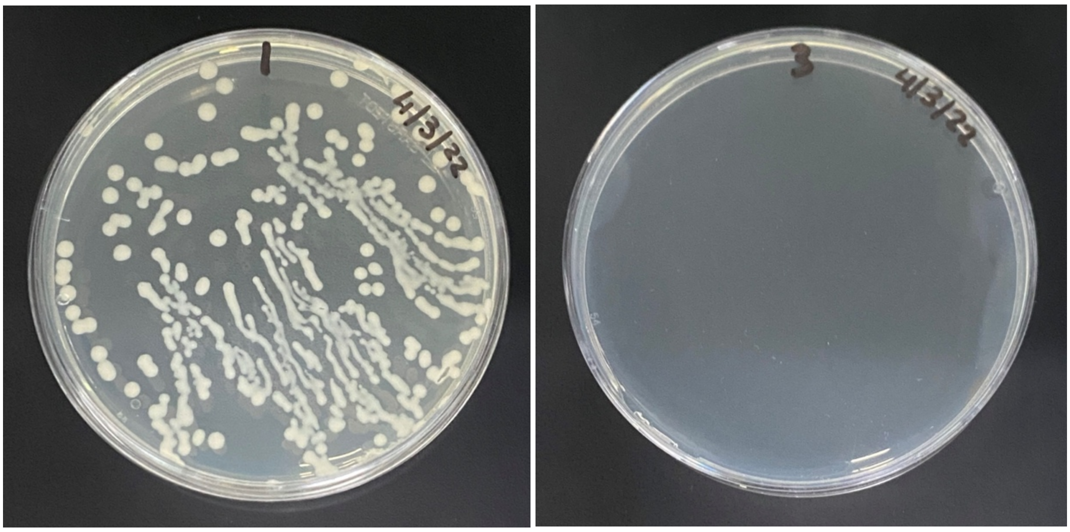


**Figure S13.** Recovery of viable colony forming units from *K. pneumoniae* ATCC 13882 cell-extracts. 100 μL of 20 mg/mL cell-extract was spread onto a nutrient agar plate before (left panel) and after filtering (right panel) and incubated at 37ºC for 16 hours.

**A** Spectinomycin

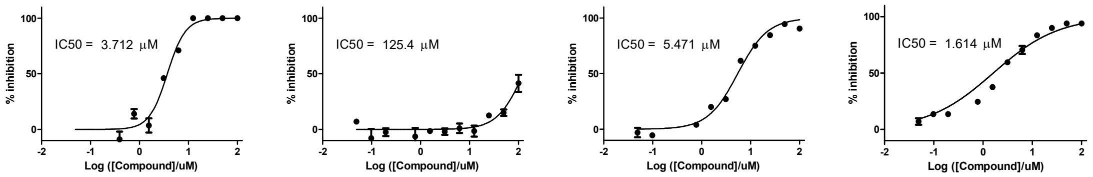


**B** Amikacin

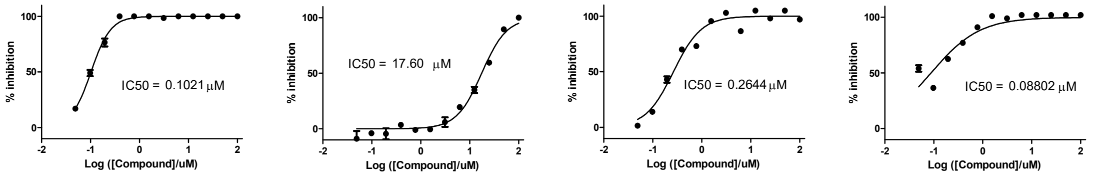


**C** Chloramphenicol

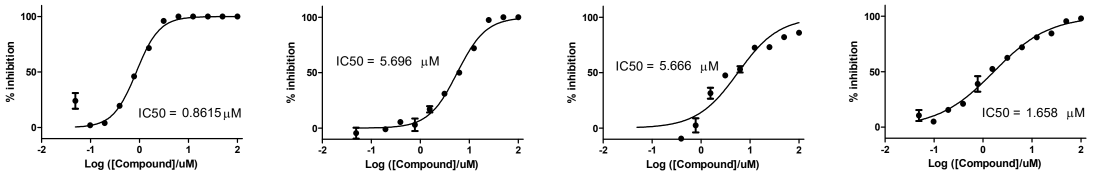


**Figure S14.** Antibiotic-resistance profiling for *K. pneumoniae* NJST258-1. Predicted (based on RESfinder) resistance was tested for spectinomycin (**A**), amikacin (**B**) and chloramphenicol (**C**). Panels (left to right) represent four biological repeats of extracts grown under standard conditions in BHI medium. Note the extracts exhibit variable resistance, likely due to absence of positive selection during cell extract processing. Data is shown as mean and standard deviation of three technical replicates.

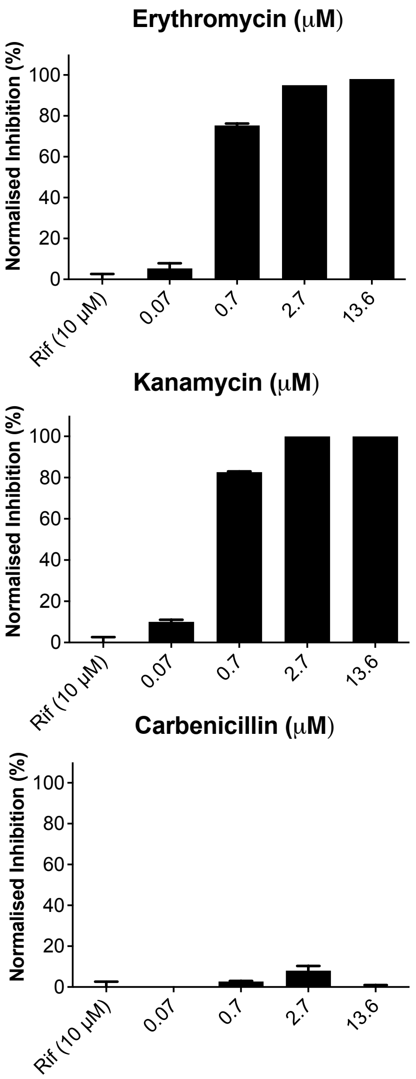


**Figure S15.** Preliminary antimicrobial inhibition of *K. pneumoniae* ATCC 13882 CFE. Data presented as normalised activity (%) for three technical repeats.

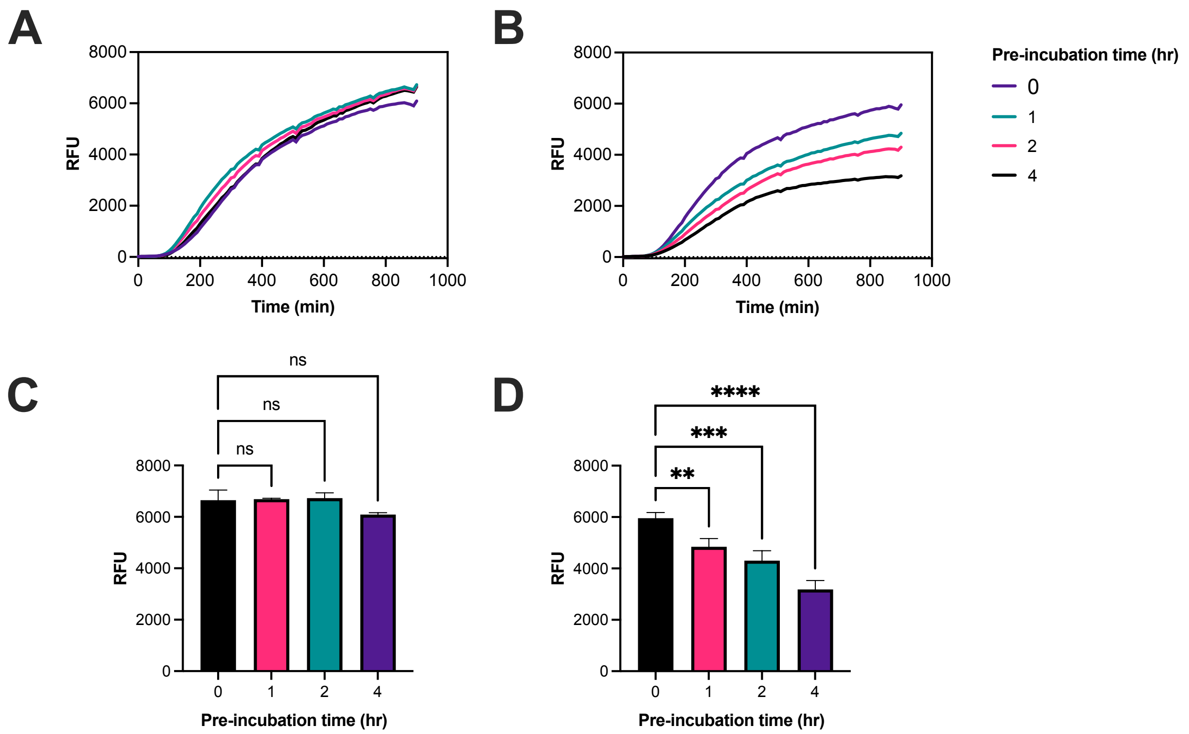


**Figure S16.** Effect of incubation time during reaction setup on assay stability. Real-time measurement of mScarlet-I fluorescence with (**A**) extracts pre-incubated at room temperature with plasmid DNA (**B**) extracts pre-incubated at room temperature with with energy mix. (**C**) End-point fluorescence data corresponding to panel A. (**D**) End-point fluorescence data corresponding to panel B. Details of experiment: After pre-incubation, the missing component of the reaction (energy mix or DNA), also incubated at room temperature, was added to complete reaction composition. The remaining protocol was followed as outlined in the methods. Standard deviation is representative of three technical measurements.

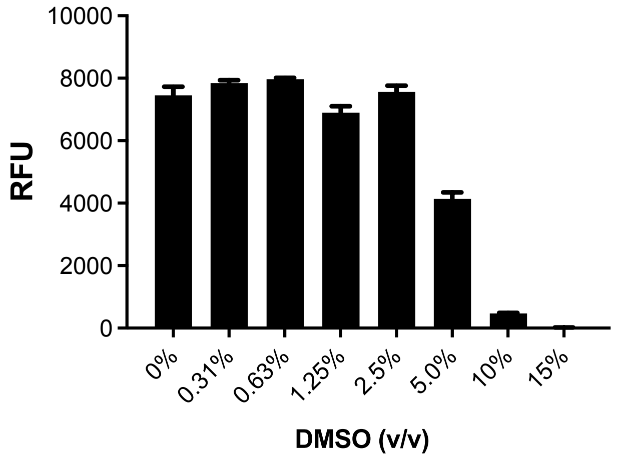


**Figure S17**. Effect of DMSO (v/v) concentration on *K. pneumoniae* ATCC 13882 CFE activity. Data is shown as mean and standard deviation of three technical measurements and is representative of two independent experiments.

**a**

| | WT | Val$^R$ |
|---|---|---|
| IC50 | 1.039 | 0.8239 |

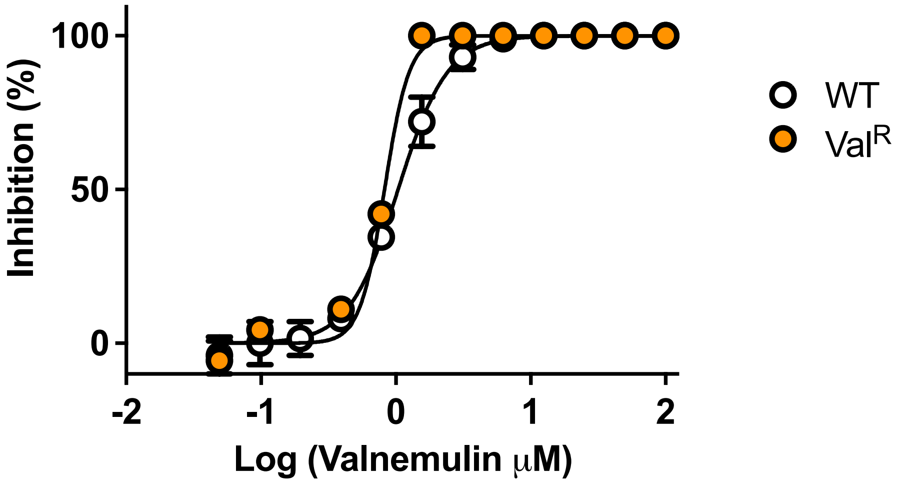


**b**

| | WT | Val$^R$ |
|---|---|---|
| IC50 | 1.039 | 0.6946 |

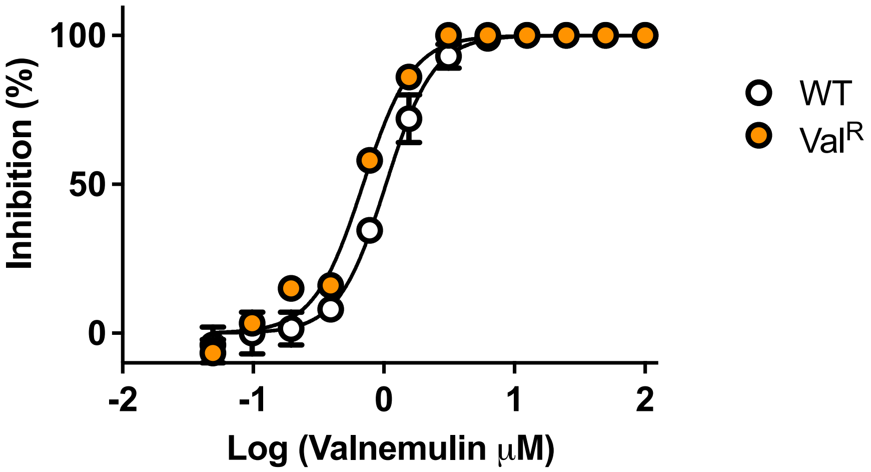


**Figure S18.** Dose-response curve of valnemulin inhibition of the ATCC 13882 (WT) and Val$^R$ CFE systems. Panels (a) and (b) represent two biological repeats, plotted separately, in comparison to a representative dataset for WT. Data for WT is shown as mean and standard error of mean of two independent experiments. See main methods for reaction setup and fluorescence settings.

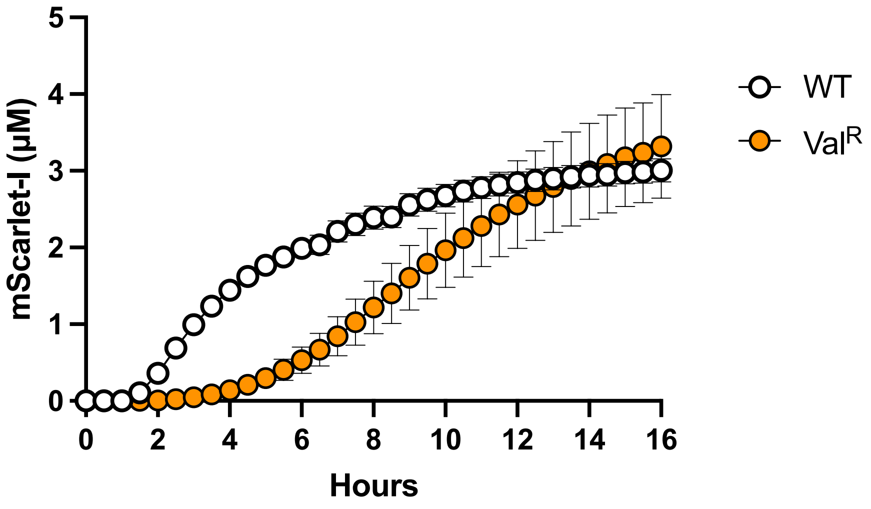


**Figure S19.** Real-time protein synthesis of the ATCC 13882 (WT) and $Val^R$ CFE systems. See main methods for reaction setup and fluorescence settings.

**Table S1 – Plasmids**

| Plasmid name | Promoter | RBS | Gene | Terminator | AddGene No. |
| --- | --- | --- | --- | --- | --- |
| Pr-deGFP-MGapt | OR2-OR1-Pr | TTTGTTTAACTTTAAGAAGGAGA | deGFP | T7 | #67734 |
| pTU1-A-SP44-mScarlet-I | SP44 | GTACTTTAACTTTAAGAAGGAGA | mScarlet-I | Bba_B0015 | #163756 |
